# Gorilla in our Midst: An online behavioral experiment builder

**DOI:** 10.1101/438242

**Authors:** Alexander Anwyl-Irvine, Jessica Massonnié, Adam Flitton, Natasha Kirkham, Jo Evershed

**Affiliations:** MRC Cognition and Brain Science Unit, University of Cambridge; Cauldron.sc: Cauldron Science, St Johns Innovation Centre, Cambridge, United Kingdom; Centre for Brain and Cognitive Development, Birkbeck College, University of London; Human Behaviour and Cultural Evolution Group, University of Exeter

**Keywords:** Online Methods, Remote Testing, Browser Timing, Attentional Control, Online Research, Timing Accuracy

## Abstract

Behavioural researchers are increasingly conducting their studies online to gain access to large and diverse samples that would be difficult to get in a laboratory environment. However, there are technical access barriers to building experiments online, and web-browsers can present problems for consistent timing – an important issue with reaction time-sensitive measures. For example, to ensure accuracy and test-retest reliability in presentation and response recording, experimenters need a working knowledge of programming languages such as JavaScript. We review some of the previous and current tools for online behavioural research, and how well they address the issues of usability and timing. We then present The Gorilla Experiment Builder (gorilla.sc) a fully tooled experiment authoring and deployment platform, designed to resolve many timing issues, and make reliable online experimentation open and accessible to a wider range of technical abilities. In order to demonstrate the platform’s aptitude for accessible, reliable and scalable research, we administered the task with a range of participant groups (primary school children and adults), settings (without supervision, at home, and under supervision, in schools and public engagement events), equipment (own computers, computer supplied by researcher), and connection types (personal internet connection, mobile phone 3G/4G). We used a simplified flanker task, taken from the Attentional Networks Task (Rueda, Posner, & Rothbart, 2004). We replicated the ‘conflict network’ effect in all these populations, demonstrating the platform’s capability to run reaction time-sensitive experiments. Unresolved limitations of running experiments online are then discussed, along with potential solutions, and some future features of the platform.

## Introduction

Behavioural research and experimental psychology are increasing their use of web-browsers and the internet to reach larger (Adjerid & Kelley, 2018), and more diverse (Casler, Bickel, & Hackett, 2013) populations than has been previously feasible with lab-based methods. However, there are unique variables which are introduced when working within an online environment. The experience of the user is the result of a large number of connected technologies. Examples of this include: the server (which hosts the experiment), the internet service provider (which delivers the data), the browser (which presents the experiment to the participant and measures their responses), and the content itself – which is a mixture of media (e.g. audio/pictures/video) and code in different programming languages (e.g. JavaScript, HTML, CSS, PHP, Java). Linking these technologies together is technically difficult, time-consuming and costly. Consequently, until recently, online research is often carried out – and scrutinized – by those with the resources to overcome these barriers.

The purpose of this paper is three-fold. Firstly, to explore the problems inherent to running behavioural experiments online with web programming languages, the issues this can create for timing accuracy, and recent improvements that can mitigate these issues. Secondly, to introduce Gorilla, an Online Experiment Builder that uses best practices to overcome these timing issues and makes reliable online experimentation accessible and transparent to the majority. Thirdly, to demonstrate the timing accuracy and reliability provided by Gorilla. This is achieved with data from a flanker task – which requires high timing fidelity - collected from a wide range of participants, settings, equipment and internet connection types.

### JavaScript

The primary consideration for online experimenters in the present time is JavaScript (JS), the language that is most commonly used to generate dynamic content on the web (such as an experiment). Its quirks (which are discussed later) can lead to problems with presentation time, and understanding it forms a large part of an access barrier.

JS is at the more dynamic end of the programming language spectrum. It is weakly typed and allows core functionality to be easily modified. Weak typing means that variables do not have declared types; the user simply declares a variable and then uses it in their code. This is in contrast to strongly typed languages, where the user must specify whether a variable they declare should be an integer, a string, or some other structure. This can lead to unnoticed idiosyncrasies – if a user writes code that attempts to divide a string by a number, or assign a number to a variable that was previously assigned to an array, JS allows this to proceed.

, JS allows users to call functions without providing all the arguments to that function. This dynamic nature gives more flexibility, but at the cost of allowing mistakes or unintended consequences to creep in. By contrast, in a strongly typed language, incorrect assignments or missing function arguments would be marked as errors that the user should correct. This results in a more brittle, but safer, editing environment. JS also allows a rare degree of modification of core structures – even the most fundamental building blocks (such as arrays) can have extra methods added to them. This can prove useful in some cases, but can easily create confusion as to which parts of the code are built-in and which parts are user defined. Together, these various factors create a programming environment that is very flexible, but one in which mistakes are easy to make and their consequences can go undetected by the designer(Richards, Lebresne, Burg, & Vitek, 2010). This is clearly not ideal for new users attempting to create controlled scientific experiments. Below we discuss two significant hurdles when building web experiments: inaccuracies in the timing of various experiment components in the browser, and the technical complexities involved in implementing an online study, including JavaScript’s contributions. These complexities present an access barrier to controlled online experiments for the average behavioural researcher.

### History of Timing Concerns

Timing concerns have been expressed regarding online studies (for an overview see Woods, Velasco, Levitan, Wan, and Spence, 2015), and while many of these concerns are historic for informed users – as solutions exist – they are still an issue for new users who may not be aware of them. Concerns fall into timing of stimuli – i.e. an image or sound is not presented for the duration you want – and the timing of response recording – i.e. the participant did not press a button at the time you think they did. These inaccuracies have obvious implications for behavioural research, especially those using time-based measures such as Reaction Time (RT).

Several things may be driving these timing issues: firstly, in JS programs, most processes within a single web-app or browser window pass through an event loop^1^ – a single thread which decides what parts of the JS code to run, and when. This loop is comprised of different queues. Queues that are managed synchronously wait until one task is complete before moving on. One example of a synchronously-managed queue is the event queue, which stores an ordered list of things waiting to be run. Queues that are managed asynchronously will start new tasks instead of waiting for the preceding tasks to finish, such as the queue that manages loading resources (e.g. images). Most presentation changes are processed through the event loop in an asynchronous queue. Thiscould be: an animation frame updating, an image being rendered, or an object being dragged around. Variance in the order in which computations are in the queue due to any experiment’s code competing with other code, can lead to inconsistent timing. When a synchronous call to the event loop requires a lot of time, it can ‘block’ the loop – preventing everything else in the queue from passing through. For instance, you may try and present auditory and visual stimuli at the same time, but they could end up out of synchronisation if blocking occurs – a common manifestation of this in web videos is unsynchronised audio and video. Secondly, the computational load on the current browser window will slow the event loop down; variance in timing is, therefore, dependent on different computers, browsers and computational load (Jia, Guo, Wang, & Zhang, 2018). For a best practices overview see Garaizar and Reips (2018).

Given the need for online research to make use of on-site computers such as in homes or in schools, the potential variance mentioned above is an important issue. A laptop with a single processor, a small amount of memory, and an out-of-date web-browser is likely to struggle to present stimuli to the same accuracy as a multi-core desktop with the most recent version of Google Chrome installed. These variances can represent variance of over 100 ms in presentation timing (Reimers & Stewart, 2016). Thirdly, by default, web browsers load external resources (such as images or videos) progressively as soon as the HTML elements that use them are added to the page. This results in the familiar effect of images ‘popping in’ as the page loads incrementally. If each trial in an online task is treated as a normal web-page, this ‘popping in’ will lead to inaccurate timing. Clearly, such a variance in display times would be unsuitable for online research, but the effect can be mitigated by loading resources in advance. A direct solution is to simply load all the required resources, for all the trials, in advance of starting the task (Garaizar and Reips, 2018). This can be adequate for shorter tasks or tasks which use a small number of stimuli, but as the loading time increases, participants can become more likely to drop out, resulting in an increase in attrition.

The same concerns (with the exception of connection speed) can be applied to the recording of response-times, which are dependent on a JS system called the ‘event system’. When a participant presses a mouse or keyboard button, recording of these responses (often through a piece of code called an ‘Event Listener’) gets added to the event loop. To give a concrete example, two computers could record different times of an identical mouse response based on their individual processing loads. It must be noted that this issue is *independent* of the browser receiving an event (such as a mouse click being polled by the operating system), where there is a relatively fixed delay, shown to be equivalent to non-browser software (de Leeuw & Motz, 2016) - this receiving delay is discussed later in the paper. Timing of event recording using the browser system clock (which some JavaScript functions do) is also another source of variance – as different machines and operating systems will have different clock accuracies and update rates.

#### Current State of the Art

Presently, the improved processing capabilities in common browsers and computers, in concert with improvements in web-language standards - such as HTML5 and ECMAScript 6 - offer the potential to overcome some concerns about presentation and response timings (Garaizar, Vadillo, & López-de Ipiña, 2012, 2014; Reimers & Stewart, 2015, 2016; Schmidt, 2001). This is because, in addition to standardised libraries (which improve the consistency of any potential web experiment between devices), these technologies use much more efficient interpreters, which are the elements of the browser which execute the code and implements computations. An example of this is Google’s V8, which improves processing speed – and therefore the speed of the event loop – significantly (Severance, 2012). In fact, several researchers have provided evidence that response times are comparable between browser-based applications and local applications (Barnhoorn, Haasnoot, Bocanegra, & Steenbergen, 2015) even in poorly standardized domestic environments - i.e. at home (Miller, Schmidt, Kirschbaum, & Enge, 2018).

A secondary benefit of recent browser improvements is scalability. If behavioural research continues to take advantage of the capacity for big-data provided by the internet, it needs to produce scalable methods of data collection. Browsers are becoming more and more consistent in the technology they adopt – meaning code will be interpreted more consistently across your experimental subjects. At the time of writing the standard for browser-based web apps is HTML5 (World Wide Web Consortium, 2019, provides current web standards) and ECMAScript JavaScript (Zaytsev, 2019, shows most browsers presently support ECMAScript 5 and above). ECMAScript (ES) is a set of standards that are implemented in JavaScript (but, can also be implemented in other environments – e.g. ActionScript in Flash), and browsers currently support a number of versions of this standard (see: Zaytsev, 2019 for details). The combination of ES and HTML5, in addition to having improved timing, is also the most scalable. They reach the greatest number of users - with most browsers supporting them, which is in contrast with other technologies, such as Java plugins and Flash that are becoming inconsistently supported – in fact Flash support has recently begun a departure from all major browsers.

### Access Barriers

Often, in order to gain accurate timing and presentation, you must have a good understanding of key browser-technologies. As in any application of computer science, there are multiple methods for achieving the same goal, and these may vary in the quality and reliability of the data they produce. One of the key resources for tutorials on web-based apps – the web itself – may lead users to use out-of-date or unsupported methods; with the fast-changing and exponential browser ecosystem, this is a problem for the average behavioural researcher (Ferdman, Minkov, Bekkerman, & Gefen, 2017). This level of complexity imposes an access barrier to creating a reliable web experiment - the researcher must have an understanding of the web ecosystem they operate in and know how to navigate its problems with appropriate tools.

There are, however, tools available which lower these barriers in various ways. Libraries, such as *jsPsych* (de Leeuw, 2015), give a toolbox of JavaScript commands which are implemented at a higher level of abstraction - therefore relieving the user of some implementation level JavaScript knowledge. Hosting tools, like ‘*Just Another Tool for Online Studies*’ (JATOS), allow users to host JavaScript and HTML studies (Lange, Kühn, & Filevich, 2015), and present these to their participants - this enables a research-specific server to be set up. However, with *JATOS* you still need to know how to set it up and manage your server, which requires a considerable level of technical knowledge. The user will also need to consider putting safeguards in place to manage unexpected server downtime caused by a whole range of issues. This may require setting up a back up system or back-up server. A common issue is too many participants accessing the server at the same time, which can cause it to overload and likely prevent access to current users mid-experiment – which can lead to data-loss (Schmidt, 2000).

The solutions above function as ‘packaged software’, where the user is responsible for all levels of implementation (i.e. browser, networking, hosting, data processing, legal compliance, regulatory compliance and insurance) - in the behavioural research use-case this requires multiple tools to be stitched together (e.g. *jsPsych* in the browser and *JATOS* for hosting). This itself presents another access barrier, as the user then must understand – to some extent – details of the web server (for instance how many concurrent connections their hosted experiment will be able to take), hosting (the download/upload speeds), the database (where and how data will be stored, e.g. in JS object notation format, or in a relational database), and how the participants are accessing their experiment and how they are connected (e.g. through *Prolific.ac* or *Mechanical Turk*).

One way to lower these barriers is to provide a platform where all of this is managed for the user commonly known as Software as a Service (SaaS) (Turner, Budgen, & Brereton, 2003). All of the above can be set up, monitored and updated for the experimenter, whilst also providing as consistent and reproducible environment as possible - something that is often a concern for web-research. One recent example of this is the online implementation of *PsyToolkit* (Stoet, 2017), where users can create, host and run experiments on a managed web server and interface - however, there is still a requirement to write out the experiment in code – representing another access limitation.

Some other tools exist in a space between SaaS and packaged software. PsychoPy3 (Peirce & MacAskill, 2018) is an open-source local application offering a graphical task builder and a python programming library. It offers the ability to export experiments built in the application (but currently not made using their python library) to JS, to a closed-source web-platform based on GitLab (an online experiment repository system) called *Pavlovia.org*, where users can host that particular task for data collection. *Lab.js* (Henninger, Mertens, Shevchenko, & Hilbig, 2017) is another task-builder, which provides a web-based GUI, where users can build a task and download a package containing the HTML, CSS and JS needed to run a study. Users are then able to export this for hosting on their own, or third-party servers. Neither of these tools function fully as SaaS, as they do not offer a fully integrated platform that allows you to build, host, distribute tasks, and manage complex experimental design (e.g. a multi-day training study) without programming, in the same environment. A full comparison of packaged software, libraries, and hosting solution can be found in Table 1.

**Table 1.**
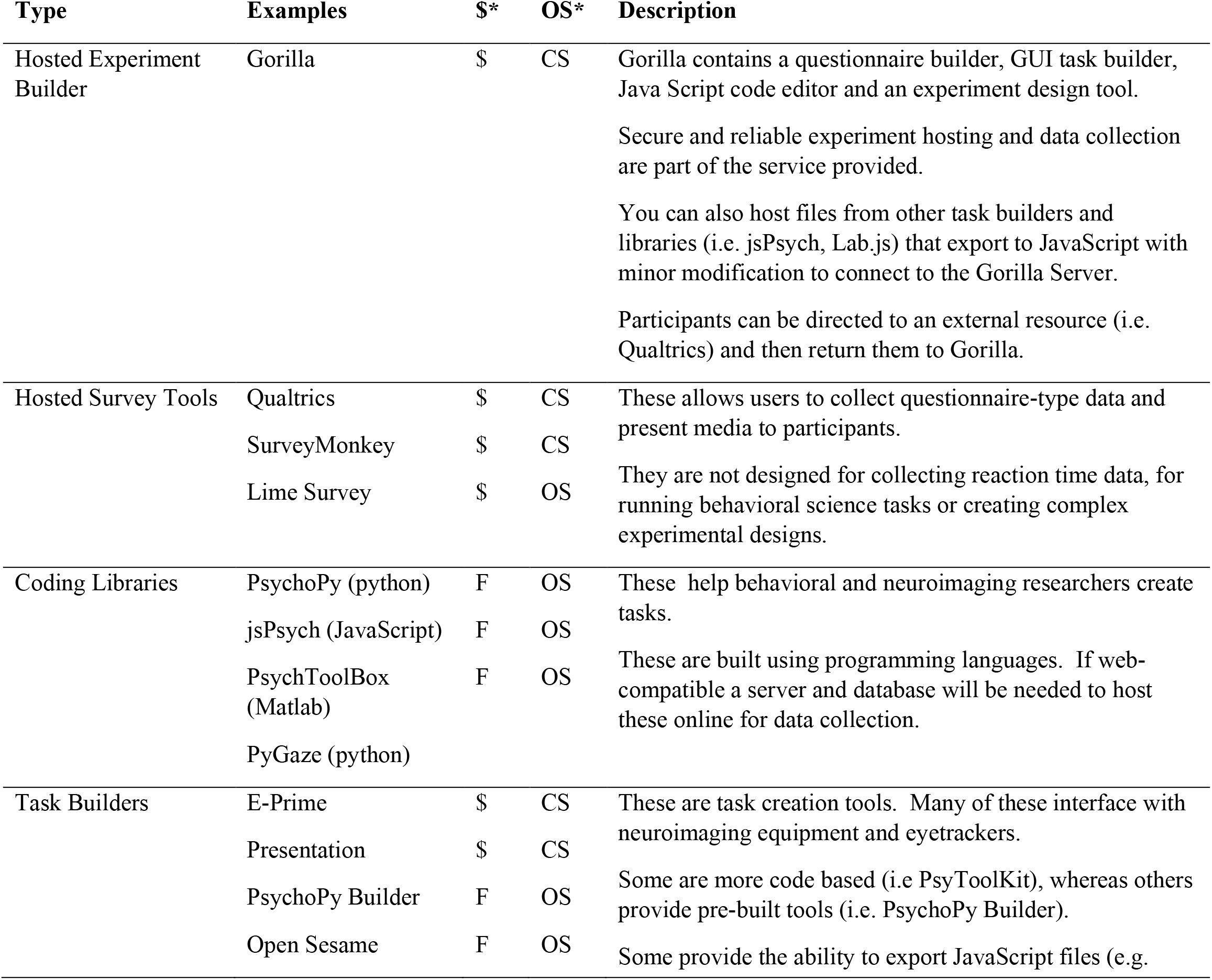
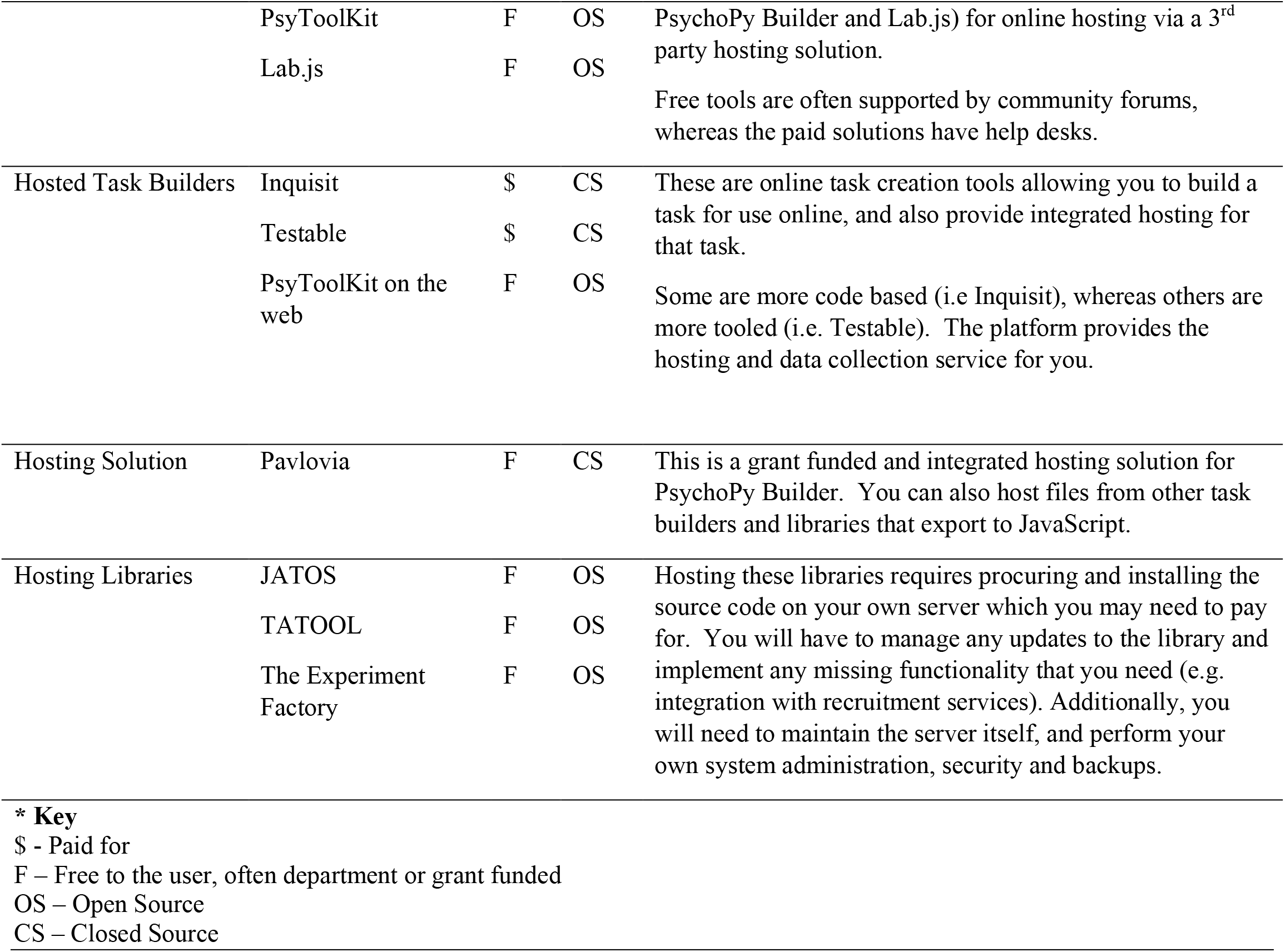
A comparison of tools available for collection of behavioural data, both online and offline.

### The Gorilla Experiment Builder

*Gorilla* (www.gorilla.sc) is an online experiment builder, and its aim is to lower the barrier to access, enabling all researchers and students to run online experiments (regardless of programming and networking knowledge). As well as giving greater access to web-based experiments, it reduces the risk of introducing higher noise in data (e.g. due to misuse of browser-based technology). By lowering the barrier, *Gorilla* aims to make online experiments available and transparent at all levels of ability. Currently, experiments have been conducted in *Gorilla* on a wide variety of topics, including: cross-lingual priming (Poort & Rodd, 2017), the provision of lifestyle advice for cancer prevention (Usher-Smith et al., 2018), semantic variables and list memory (Pollock, 2018), narrative engagement (Richardson et al., 2018), trust and reputation in the sharing economy (Zloteanu, Harvey, Tuckett, & Livan, 2018), how individual’s voice identities are formed (Lavan, Knight, & McGettigan, 2018), and auditory perception with degenerated music and speech (Jasmin, Dick, Holt, & Tierney, 2018). Also, several studies have pre-registered reports, including: object size and mental simulation of orientation (Chen, de Koning, & Zwaan, 2018) and the use of face regression models to study social perception (Jones, 2018). Additionally, It has also been mentioned in a paper on the gamification of cognitive tests (Lumsden, Skinner, Coyle, Lawrence, & Munafo, 2017). Gorilla was launched in September 2016, and as of January 2019 there are over 5,000 users signed up to Gorilla, across more than 400 academic institutions. In the last three months of 2018, data was collected from over 28,000 participants – an average of around 300 participants per day.

One of the largest differences between Gorilla and the other tools mentioned above (a comprehensive comparison of these is in Table 1) is that it is an *experiment design tool*, not just a task building or questionnaire tool. At the core of this is the Experiment Builder, a graphical tool which allow you to creatively reconfigure task and questionnaires into a wide number of different experiment designs without having to code. The interface is built around dragging and dropping nodes (which represent what the participant sees at that point, or modifications to their path through the experiment) and connecting them together with arrow lines. This modular approach makes it much easier for labs to reuse elements that have been created before by themselves or by others. For instance, this allows any user to construct complex counterbalanced, randomised, between-subjects designs with multi-day delays and email reminders with absolutely no programming needed. Examples of this can be seen in Table 2.

**Table 2.**
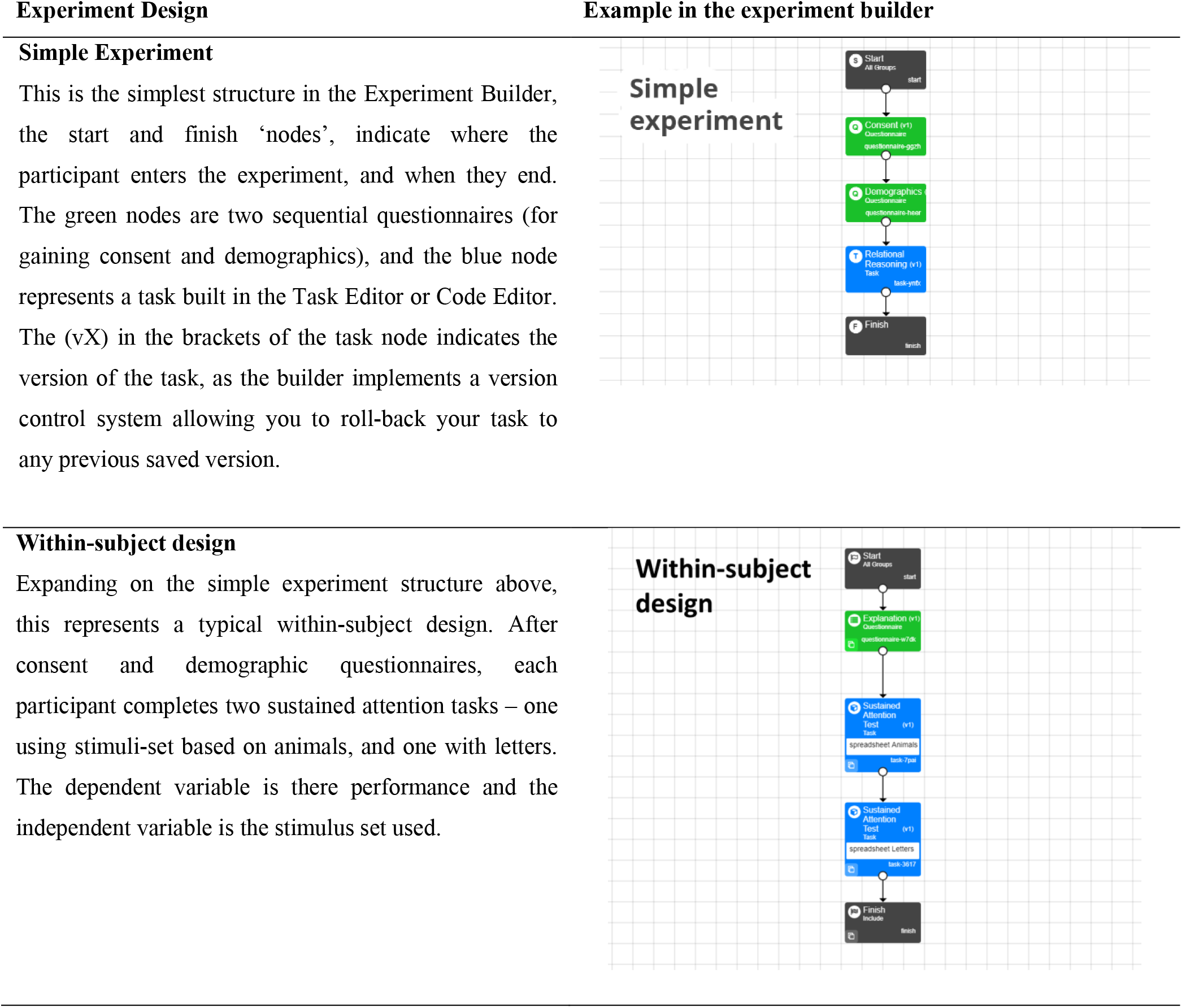
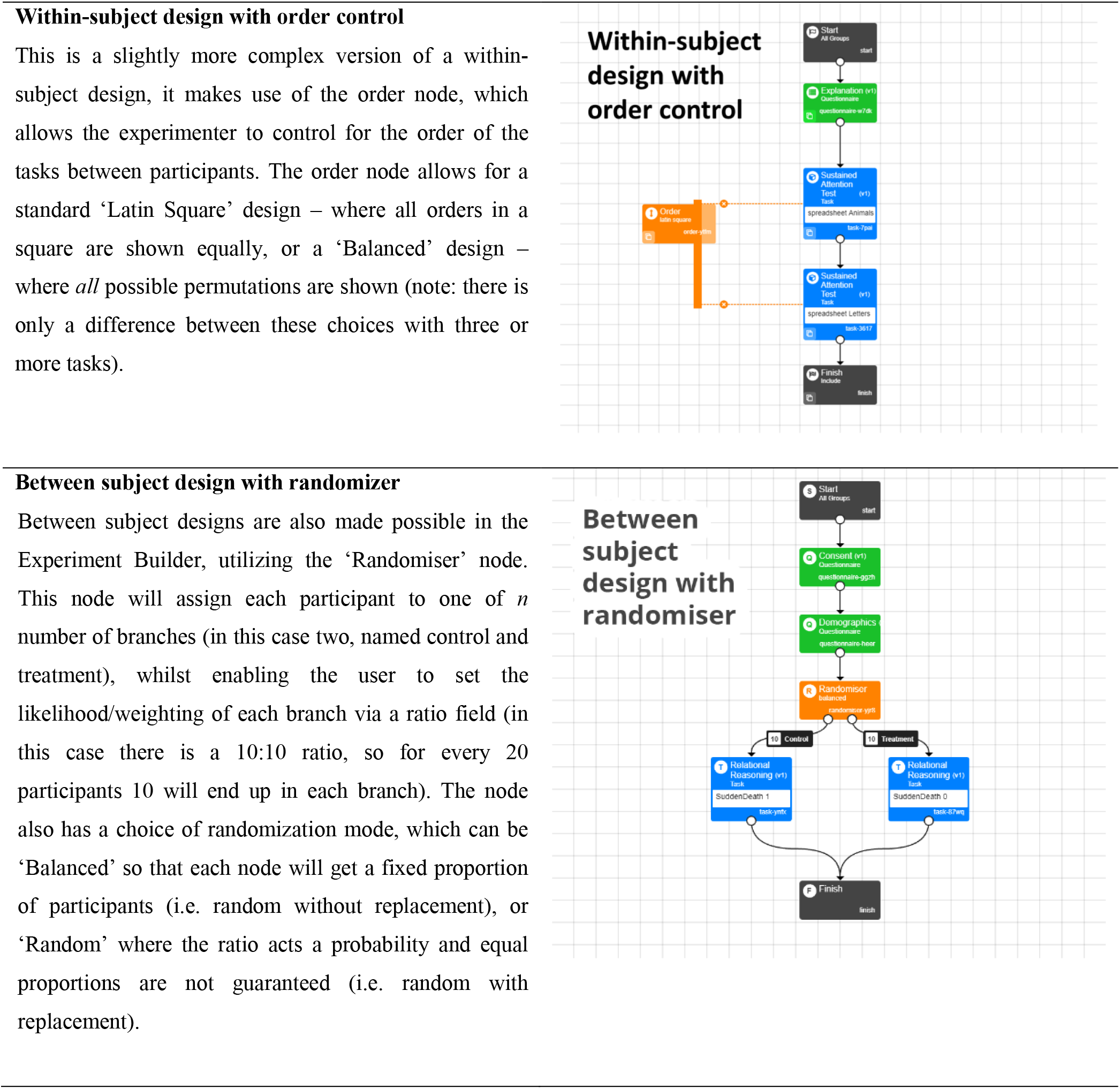
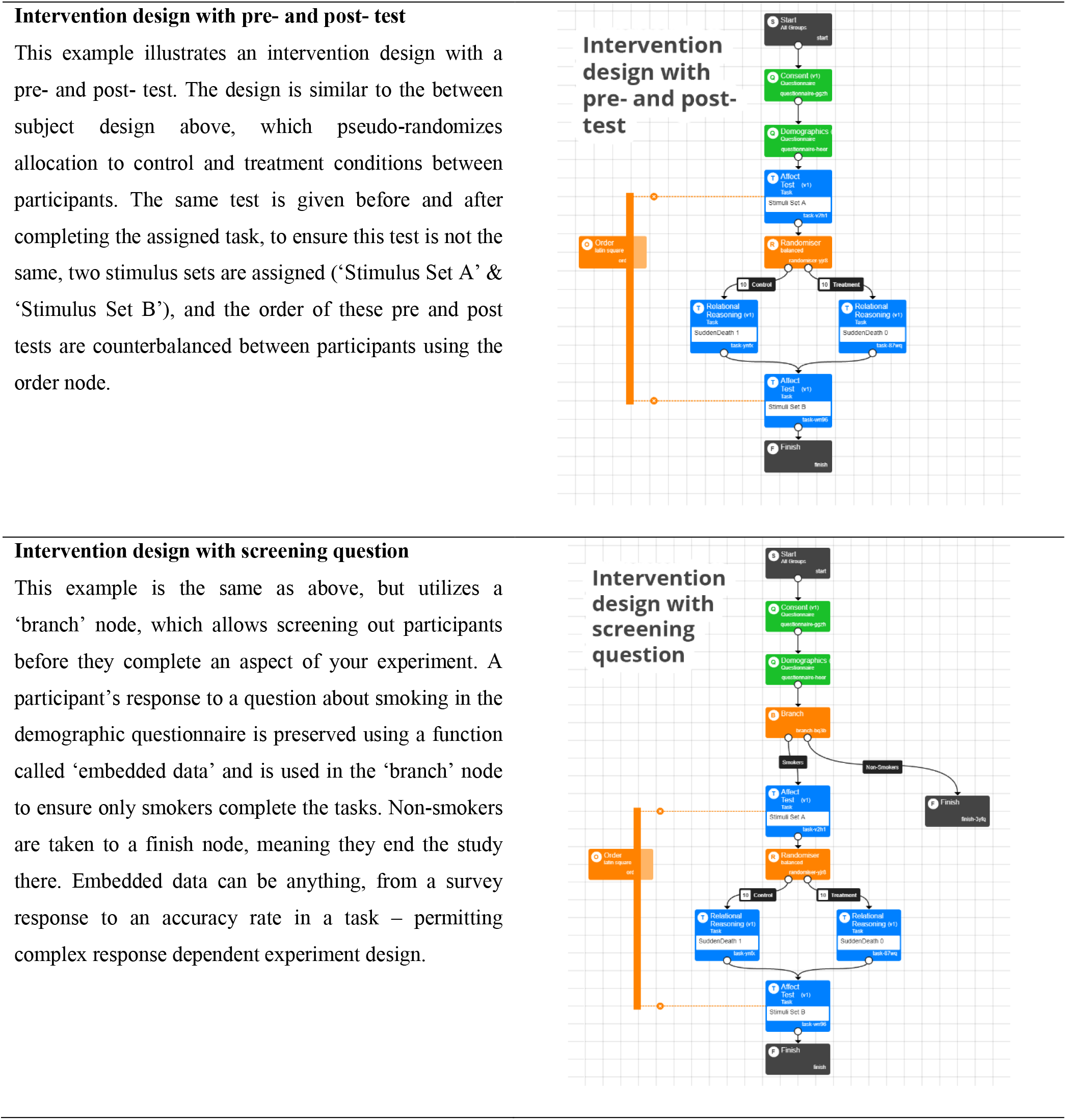
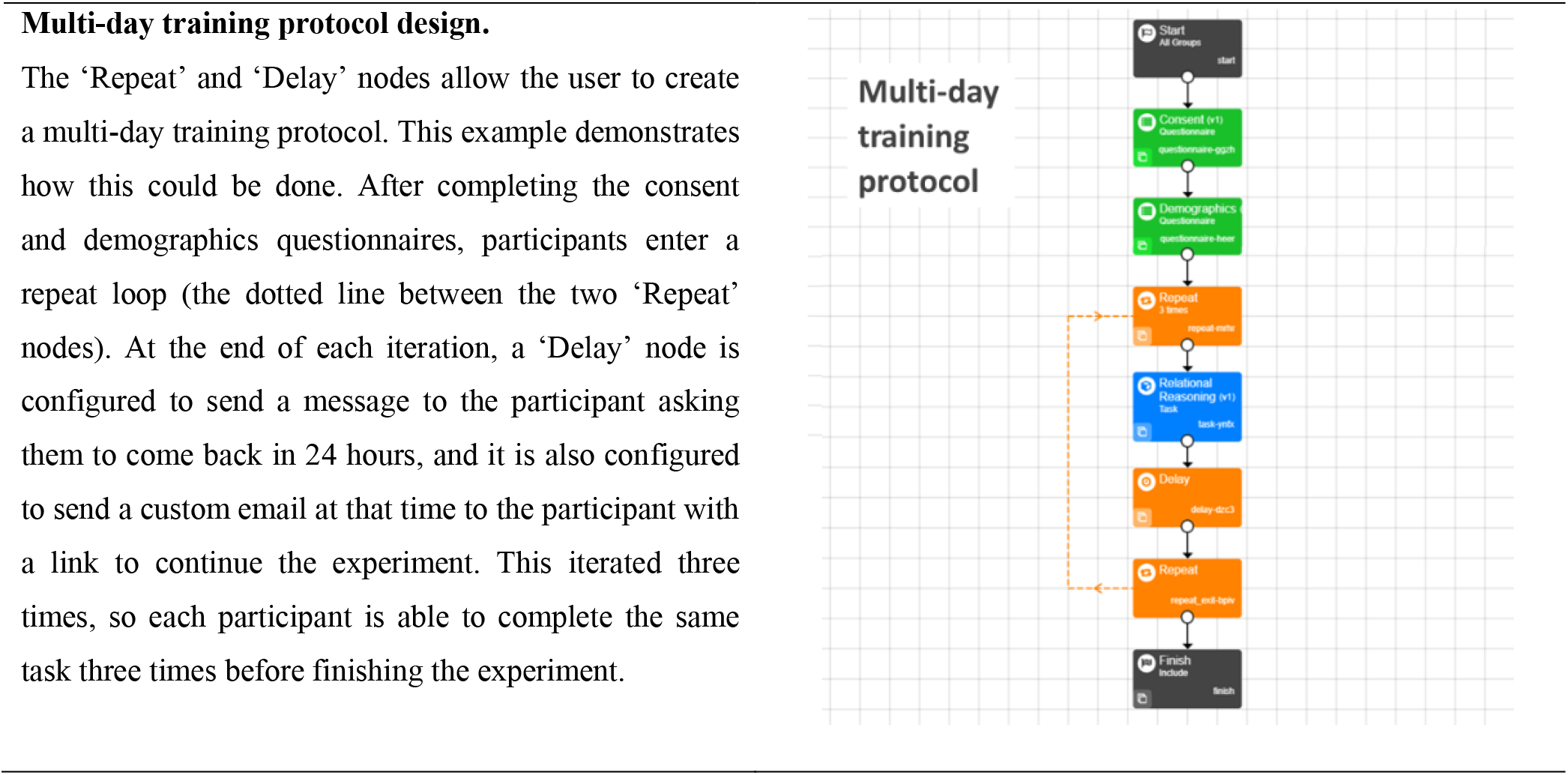
Examples of experiment designs possible within Gorilla’s Experiment Builder interface.

*Gorilla* provides researchers with a managed environment in which to design, host and run experiments. It is fully compliant with the EU General Data Protection Regulation (GDPR), and NIHR and BPS guidelines, and has backup communication methods for data in the event of server problems (to avoid data loss). A graphic user interface (GUI) is available for building questionnaires (called the ‘Questionnaire Builder’), experimental tasks (the ‘Task Builder’) and running the logic of experiments (the ‘Experiment Builder’). For instance, a series of different attention and memory tasks could be constructed with the Task Builder, and then their order of presentation is controlled with the Experiment Builder. Both are fully implemented within a web-browser and are illustrated in Figure 1. This allows users with little or no programming experience to run online experiments, whilst controlling and monitoring presentation and response timing.

**Fig. 1:**
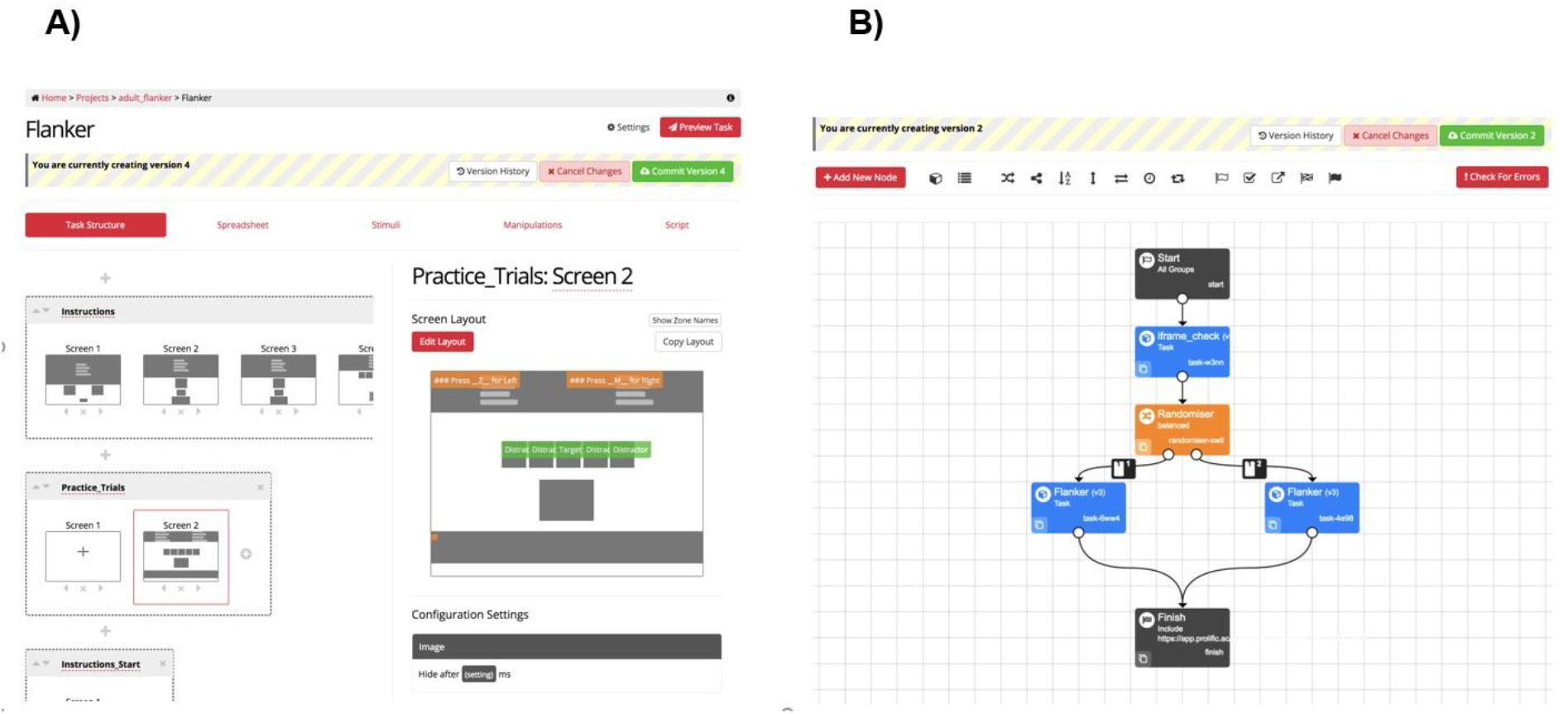
An example of the two main GUI elements of Gorilla. A) shows the task builder, with a screen selected, showing how a trial is laid out B) shows the experiment builder, there is a check for the participant, followed by a randomiser node which allocates them to one of two conditions, before sending them to a Finish node.

At the Experiment Builder level (Figure 1B) users can create logic for the experiment though it’snodes that manage capabilities such as: randomisation, counterbalancing, branching, task switching, repeating and delay functions. This range of functions makes it as easy to create longitudinal studies with complex behavior. An example could be a 4 week training study with email reminders, where participants receive different tasks based on prior performance, or the experiment tree just as easily enables a one-shot between subject experiment. Additionally, *Gorilla* includes a redirect node that allows users to redirect participants to another hosted service and then send them back again. This allows users to use the powerful Experiment Builder functionality (i.e. multi-day testing) while using a different service (such as *Qualtrics*) at the task or questionnaire level. Table 2 provides a more detailed explanation of several example experiments made in the builder.

The Task Builder (Figure 1A) provides functionality at the task level. Each experimental task is separated into ‘displays’ that are made of a sequence of ‘screens’. Each screen can be configured by the user to contain an element of a trial, be that: text, images, videos, audio, buttons, sliders, keyboard responses, progress bars, feedback and a wide range of other stimuli and response options. See the full list here: https://gorilla.sc/support/articles/features. The content of these areas can be either static (such as instructions text), or change on a per-trial basis (where the content is set using a spreadsheet). The presentation order of these screens is dependent on sequences defined in this same spreadsheet, where blocked or complete randomisation can take place on the trial level. Additionally, the Task Builder also has a ‘Script’ tab, which allows the user to augment the functionality provided by Gorilla with JS. This allows users to use the GUI and JS side-by-side. There is also a separate ‘Code Editor’ which provides a developmental environment to make experiments purely in code. This allows users to include external libraries – such as jsPsych. The purpose of the Code Editor is to provide a secure and reliable service for hosting, data storage and participant management for tasks written in code.

Using tools like the ‘Code Editor’, users can extend the functionality of *Gorilla* through use of the scripting tools, where custom JavaScript commands, HTML templates and an application programming interface (API) are available – an API is a set of functions which gives access to the platforms functionality in the code editor, and also allows users integrate 3^rd^ party libraries into their experiments (tasks programmed in jsPsych for instance). Therefore, *Gorilla* also can function as a learning platform where users progress on to programming - whilst providing an API that manages more complex issues (such as timing and data management) where a beginner might make errors. The code editor allows inclusion of any external libraries (e.g. animation: *pixi.js,* image processing: *OpenCV.js,* eyetracking: *WebGazer.js*).. A full list of features is available here: www.gorilla.sc/tools, and a tutorial is included in the supplementary materials below.

#### Timing Control

A few techniques are utilised within *Gorilla* to control timing. To minimise any potential delays due to network speed (mentioned above), the resources from several trials are loaded in advance of presentation, a process called caching. Gorilla loads the assets required for the next few trials, begins the task, and then continues to load assets required for future trials while the participant completes the task. This strikes an optimal balance between ensuring that trials are ready to be displayed when they are reached, while preventing a lengthy load at the beginning of the task. This means that fluctuations in connection speed will not lead to erroneous presentation times. The presentation of stimuli are achieved using the requestAnimationFrame() function, which allows the software to count frames and run code when the screen is about to be refreshed ensuring screen-refreshing in the animation loop does not cause hugely inconsistent presentation. This method has previously been implemented to achieve accurate audio presentation (Reimers & Stewart, 2016) and accurate visual presentation (Yung, Cardoso-Leite, Dale, Bavelier, & Green, 2015). Rather than assuming that each frame is going to be presented for 16.667 ms, and presenting a stimulus for the nearest number of frames (something that commonly happens), *Gorilla* times each frame’s actual duration - using requestAnimationFrame(). The number of frames a stimulus is presented for can, therefore, be adjusted depending on the duration of each frame - so that most of the time a longer frame refresh (due to lag) will not lead to a longer stimulus duration. This method was used in the (now defunct) *QRTEngine* (Barnhoorn et al., 2015), and to our knowledge is not used in other experiment builders (for a detailed discussion on this particular issue see this GitHub issue: www.github.com/jspsych/jsPsych/issues/75 and this blog post on the *QRTEngine*’s website: www.qrtengine.com/comparing-qrtengine-and-jspsych/).

Reaction time (RT) is measured, and presentation time recorded using the performance.now() function, which is independent of the browser’s system clock, and therefore not impacted by changes to this over time. This is the same method used by *QRTEngine*, validated using a photodiode (Barnhoorn et al.,2015). Whilst performance.now(), and associated high resolution time-stamps, offer the greatest accuracy, the resolution has been reduced intentionally by all major browsers to mitigate against certain security threats (Schwarz, Maurice, Gruss & Mangard, 2017; Kocher et al., 2018). In most browsers the adjusted resolution is rounded to the nearest 1-5 ms, with 1 ms being the most common (Mozilla, 2019) – this is unlikely to be a permanent change, and will be improved when the vulnerabilities are better understood (Ritter, Mozilla, 2018; Mozilla, 2019).

Additionally, to maximise data quality, the user can restrict through the GUI which devices, browsers and connection speed they will allow the participant to have, and all this data is then recorded. This method allows the restriction of the participant environment, where only modern browser/device combinations are permitted - so the above techniques - and timing accuracy - are enforced. The user is able to make their own call in a trade-off between potential populations of participants, and restrictions on them to promote accurate timing, dependent on the particulars of the task or study.

### Case Study

As a case study, an experiment was chosen to illustrate the platform’s capability for accurate presentation and response timing. To demonstrate *Gorilla*’s ability to work within varied setups, different participant groups (primary school children and adults in both the UK and France), settings (without supervision, at home, and under supervision, in schools and in public engagement events), equipment (own computers, computer supplied by researcher), and connection types (personal internet connection, mobile phone 3G/4G) were selected.

We ran a simplified flanker task taken from the Attentional Networks Task (ANT) (Fan, McCandliss, Sommer, Raz, & Posner, 2002; Rueda et al., 2004). This task measures attentional skills, following the Attentional Network theory. In the original ANT papers, three attentional networks are characterised: alerting (a global increase in attention, delimited in time but not in space), orienting (the capacity to spatially shift attention to an external cue), and executive control (the resolution of conflicts between different stimuli). For the purpose of this paper and for the sake of simplicity we will focus on the “Executive control” component. This contrast was chosen as MacLeod et al. (2010) found that it was highly powered and reliable relative to the other conditions in the ANT. Participants responded as quickly as possible to a central stimulus, one that is either pointing in the same direction as identical flanking stimuli, or in the opposite direction. Thus, there are both congruent (same direction) and incongruent (opposite direction) trials.

Research with this paradigm robustly shows that RTs to congruent trials are faster than those to incongruent trials – Rueda et al. (2004) term this the ‘conflict network’. This RT difference, while significant, is often less that 100 ms, and thus very accurately timed visual presentation, and accurate recording of responses is necessary. Crump, McDonnell, and Gureckis (2013) successfully replicated the results of a similar flanker task online, using Mechanical Turk, with letters as targets and flankers, so we know this can be an RT-sensitive task that works online. Crump et al. (2013) coded this task in JavaScript and HTML and managed the hosting and data-storage themselves; however, this current version was created and run entirely using *Gorilla*’s GUI. It is hypothesised that the previously recorded conflict RT difference will be replicated on this platform.

## Experiment 1

### Methods

#### Participants

Data was drawn from three independent groups. Group A was in Corsica, France, across 6 different primary classrooms. Group B was in three primary schools in London, UK. Group C was at a public engagement event carried out at a university in London.

In total, 270 elementary school children were recruited. Two participants were excluded for not performing above chance (< 60% accuracy) in the task. The final sample included 268 children (53.7% of females), between 4.38 and 12.14 years of age (M = 9.21; SD = 1.58). Details about the demographics for each group are provided in Table 3. Informed written parental consent was obtained for each participant, in accordance with the University’s Ethics Committee.

**Table 3:** Sample Size, Age and Gender of the participants for each of the three groups. Age Range represented by Min and Max columns. Group A was children in school in Corsica, France, Group B consisted of children in schools in London, UK, Group C consisted of children attending a university public engagement event in London.

|  | *Size* | *Gender (% female)* | *Age* |  |  |  |
| --- | --- | --- | --- | --- | --- | --- |
|  |  |  | *Min* | *Max* | *Mean* | *SD* |
| **Group A** | 116 | 49.1 | 7.98 | 11.38 | 9.95 | .69 |
| **Group B** | 43 | 60.5 | 8.82 | 11.19 | 9.85 | .55 |
| **Group C** | 109 | 56.0 | 4.38 | 12.14 | 8.18 | 1.93 |

#### Procedure

In all three groups, participants were tested in individual sessions, supervised by a trained experimenter. Although great care was taken to perform the task in a quiet place, noise from adjacent rooms sometimes occurred in the school groups (A and B). To prevent children from getting distracted, they were provided with noise cancelling headphones (Noise Reduction Rating of 34dB; ANSI S3.19 and CE EN352-1 Approved).

The task was carried out using the web-browser Safari, on a Mac OS X operating system. Because a stable internet connection was often lacking in schools, in groups A and B, a mobile phone internet connection was used – this could vary from 3G to 4G.

#### Flanker Task

The flanker task was adapted from Rueda et al. (2004). A horizontal row of five cartoon fish were presented in the centre of the screen (see Figure 2), and participants had to indicate the direction the middle fish was pointing (either to the left, or right), by pressing the “X” or “M” buttons on the keyboard. These buttons were selected so that children could put one hand on each response key. Buttons were covered by arrows stickers (left arrow for “X”; right arrow for “M”) to avoid memory load. The task has two trial types: *congruent* and *incongruent*. In *congruent* trials, the middle fish was pointing in the same direction as the flanking fish. In the *incongruent* trials, the middle fish was pointing in the opposite direction. Participants were asked to answer as quickly and accurately as possible.

**Fig. 2:**
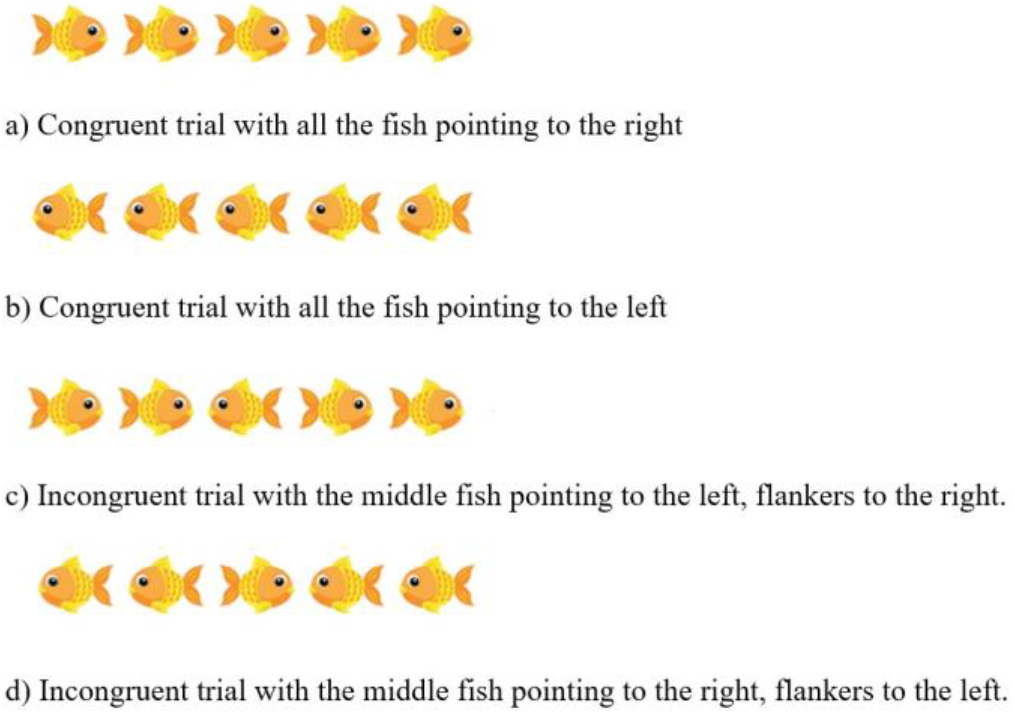
Trial types for Experiment 1. The different conditions used in the flanker task.

After the experimenter had introduced the task, there were 12 practice trials, with immediate feedback on the screen. A red cross was displayed if children answered incorrectly, and a green tick was shown if they answered correctly. Instructions were clarified by the experimenter if necessary. After the practice trials, four blocks of 24 trials each were presented. Self-paced breaks were provided between the blocks. For each participant, 50% of the trials were congruent, and the direction of the middle fish varied randomly between left and right. Four types of trials were therefore presented (see Figure 2): all the fish pointing to the right (25%), all the fish pointing to the left (25%), middle fish pointing to the right and flanking fish to the left (25%), middle fish pointing to the left and flanking fish to the right (25%).

As shown in Figure 3, for each trial, a fixation cross was displayed for 1700 ms. The cross was followed by the presentation of the fish stimuli, which stayed on screen until a valid response (either “X” or “M”) was provided. A blank screen was then displayed before the next trial. The duration of the blank screen varied randomly between 400, 600, 800 and 1000 ms. Overall, the task took no more than 10 minutes.

**Fig. 3:**
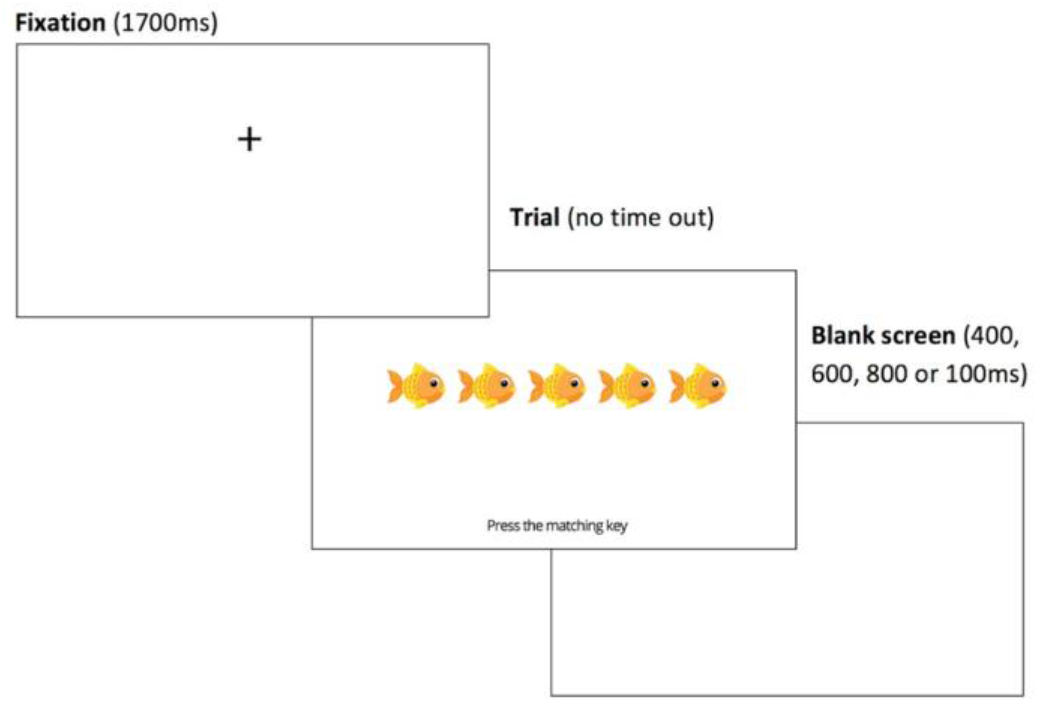
The time-course of a typical trial in Experiment 1. These screens represent what the participant was seeing within their web-browser.

#### Power Calculations

The main flanker effect reported in Rueda et al. (2004)’s ANT ANOVA results (Experiment 1) was F(2, 88) = 61.92; *p* = <0.001. They did not report the effect size, so this permits us only to estimate the effect size using partial eta squared. This was calculated using the calculator provided by Lakens (2013), as *η*_p^2^_ = .58 (95% CI= .44 - .67). Using G*power (Faul, Erdfelder, Buchner, & Lang, 2009), an a priori power calculation was computed for a mixed-factor ANCOVA with 3 groups, and a measurement correlation (congruent*incongruent) of .81 (taken from internal correlation of this measure reported in MacLeod et al. (2010)). In order to reach a power of above .95 a sample of 15 would be needed for each of our groups - we included in excess of this to increase sensitivity and provide power of <.99.

### Results

**Table 4:**
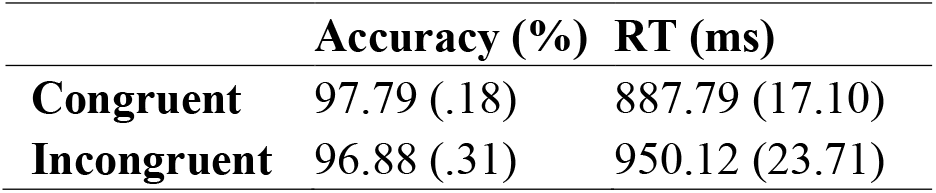
Accuracy and Reaction Time of participants, averaged (mean) over all groups, split by Congruency. Standard Error of the means are shown in brackets.

#### Data pre-processing

Reactions Time for correct answers (RTs) were computed. RTs under 200 ms were excluded, being too short to follow the perception and generation of response to the stimulus, and therefore are likely to be the result of anticipatory guesses, and do not relate to the process of interest (Whelan, 2008, also see studies on visual awareness from Koivisto & Grassini, 2016 and Rutiku, Aru, Bachmann, 2016).

Furthermore, RTs above 3 standard deviations from the mean of each subject were excluded in order to prevent extreme values from influencing the results (in some instances, children were asking a question in the middle of the trial) (Whelan, 2008).

The accuracy score (number of correct answers / total number of trials) was calculated after trials were excluded for having a RT greater than three standard deviations from the mean, and/or less than 200ms.

#### Accuracy

A mixed-factor ANCOVA was performed, with Congruency as a within-subject factor (2 levels: Accuracy for congruent trials, Accuracy for incongruent trials), Group as a between-subject factor (3 levels: Group A, Group B, Group C), and Age as a covariate. There was a significant main effect of Congruency on participants’ accuracy (*F* (1, 264) = 9.02, *p* = .003, *η*_p^2^_ = .033). Although performance was at ceiling for both types of trials, participants were more accurate for congruent trials, compared to incongruent trials (see Table 2). This effect significantly interacted with participants’ age, (*F* (1, 264) = 6.80, *p* = .010, *η*_p^2^_ = .025), but not with participants’ group (*F* (2, 264) = .501, *p* = .607, *η*_p^2^_ = .004). In order to shed light on this interaction effect, the difference in accuracy scores between congruent trials and incongruent trials was computed for each subject. This difference diminished with age (*r* = -.22; *p* < .001).

Results from the ANCOVA should however be interpreted with caution, since two assumptions were violated with the present data. First, the distribution of accuracy scores in each of the three groups were skewed and did not follow a normal distribution (for Group A: Shapiro-Wilk *W* = .896, *p* < .001; for Group B, *W* = .943, *p* = .034; for Group C: W = .694, *p* < .001). Secondly, the Levene’s test for equality of variance between groups was significant (for Congruent trials: *F*(2, 265) = 5.75, *p* = .004; for Incongruent trials: *F*(2, 265) = 13.90, *p* < .001). The distribution of data is represented in Figure 4.

**Fig. 4:**
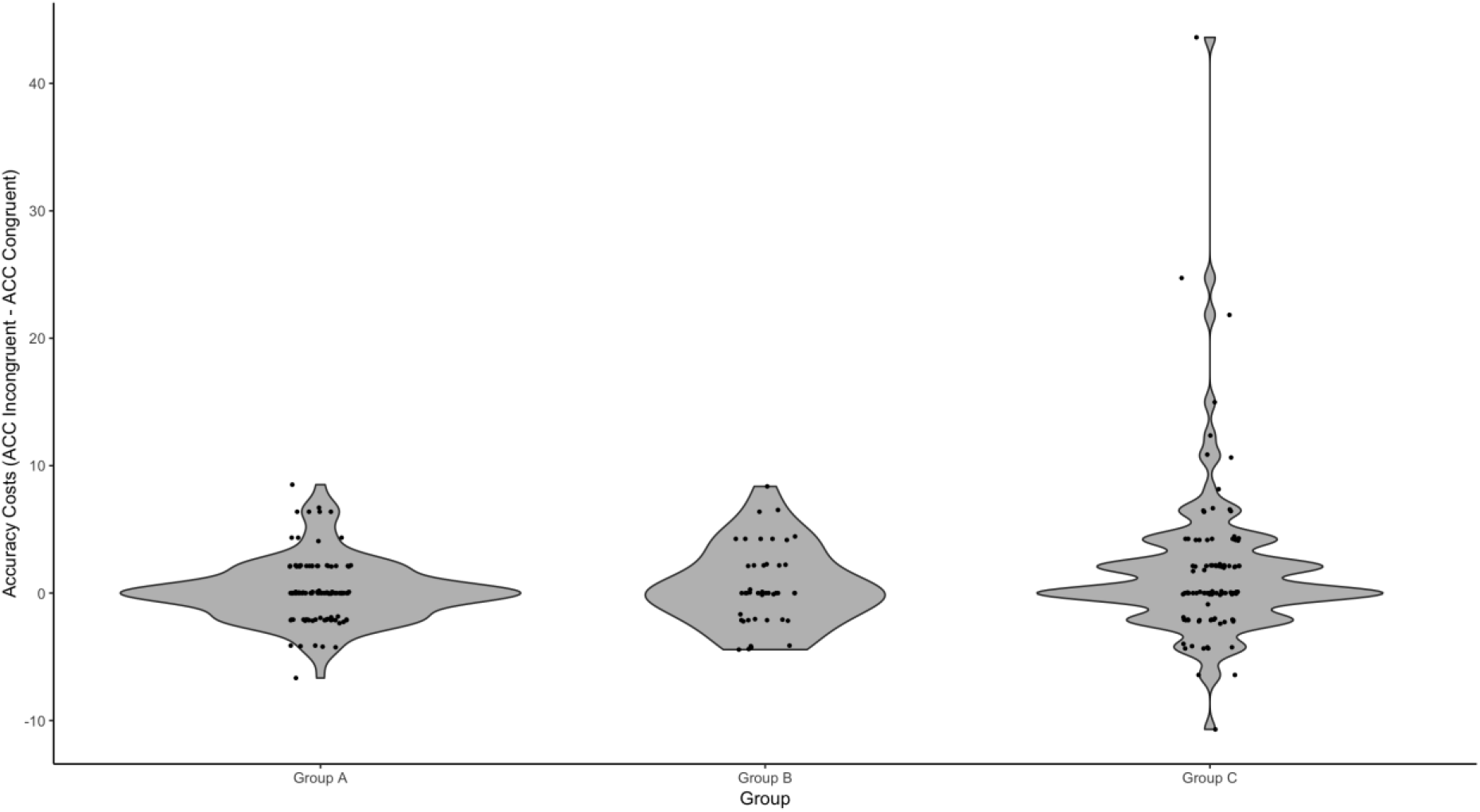
Distribution of Accuracy difference between congruent and incongruent trials, for each group in Experiment 1. Group A was children in school in Corsica, France, Group B consisted of children in schools in London, UK, Group C consisted of children attending a university public engagement event in London.

Due to these violations, the non-parametric Friedman Test was carried out, which is tolerant of non-normality, and it also revealed a significant effect of Congruency on accuracy scores (*χ*^2^(1) = 5.17, *p* < .023). Further non-parametric tests were also carried out to test whether the congruency effect differed between the three groups of participants. A Welch test for independent samples, tolerant of the non-equal variances between groups, indicated that the difference in accuracy between congruent and incongruent trials was not similar across groups (*F*(2, 108.53) = 3.25, *p* = .042). Games-Howell post hoc comparisons indicated that is effect was driven by the difference between Group A and Group C (*p* = .032). Group A and Group B did not significantly differ from each other (*p* = .80), neither did Group B and Group C (*p* = .21). Descriptive statistics are reported in Table 5.

**Table 5:** Average difference in accuracy between congruent and incongruent trials, per participants’ group. Standard Error of the means are shown in brackets.

|  | Accuracy Difference<br>(Accuracy Congruent – Accuracy Incongruent) |
| --- | --- |
| Group A | .18 (.23) |
| Group B | .52 (.47) |
| Group C | 1.82 (.60) |

However, as reported in Table 3, participants in Group C were younger than in Group A, and the difference in accuracy between congruent and incongruent trials is larger for younger children. In order to check if the group differences revealed by the Welch test were driven by this age difference, the ‘Accuracy Difference’ scores (Congruent – Incongruent Accuracy) were regressed on Age, and the Welch test was performed on the residuals. The difference between participants’ group was then non-significant (*F*(2, 108.48) = .396, *p* = .674), indicating the previous Welch test results were likely driven by age.

#### Reaction Time

A mixed-factor ANCOVA was performed, with Congruency as a within-subject factor (2 levels: RT for congruent trials, RT for incongruent trials), Group as a between-subject factor (3 levels: Group A, Group B, Group C), and Age as a covariate. There was a main effect of Congruency on participants’ RTs (*F* (1, 264) = 18.92, *p* < .001, n^2^p = .067). Participants took longer to provide the correct answer for incongruent trials, compared to congruent trials (see Table 2). This effect significantly interacted with Age, (*F* (1, 264) = 11.36, *p* = .001, n^2^p = .041), but not with Group type (*F* (2, 264) = .594, *p* = .553, n^2^p = .004). In order to better understand this interaction effect, RTs costs were calculated by subtracting the mean RTs to the congruent trials, to the mean RTs to incongruent trials. Higher values indicate poorer inhibitory control, in that it takes longer to give the correct answer for incongruent trials. RTs costs decreased with Age, indicating an improvement in inhibitory control over development (r = -.20; *p* = .001).

Similarly to the analyses for accuracy scores, RTs in each of the three groups were skewed and do not follow a normal distribution (for Group A: Shapiro-Wilk W = .476, *p* < .001; for Group B, W = .888, *p* = .034; for Group C: W = .649, *p* < .001). Secondly, the Levene’s test for equality of variance between groups was significant (for Congruent trials: *F*(2, 265) = 9.36, *p* < .001; for Incongruent trials: *F*(2, 265) = 7.28, *p* < .001). The distribution of data is represented in Figure 5. The non-parametric Friedman Test, which is tolerant to non-normal data, also reveals a significant effect of Congruency on RTs for correct answers (X^2^ (1) = 55.37, *p* < .001). A non-parametric Welch test for independent samples – tolerant of the non-equal distributions between groups - was carried out, indicating that RT costs (difference between congruent and incongruent trials) did not significantly differ between the three groups of participants (*F*(2, 165.22) = .335, *p* = .716), indicating that the main effect in the ANCOVA was unlikely to be driven by the group’s differences.

**Fig. 5:**
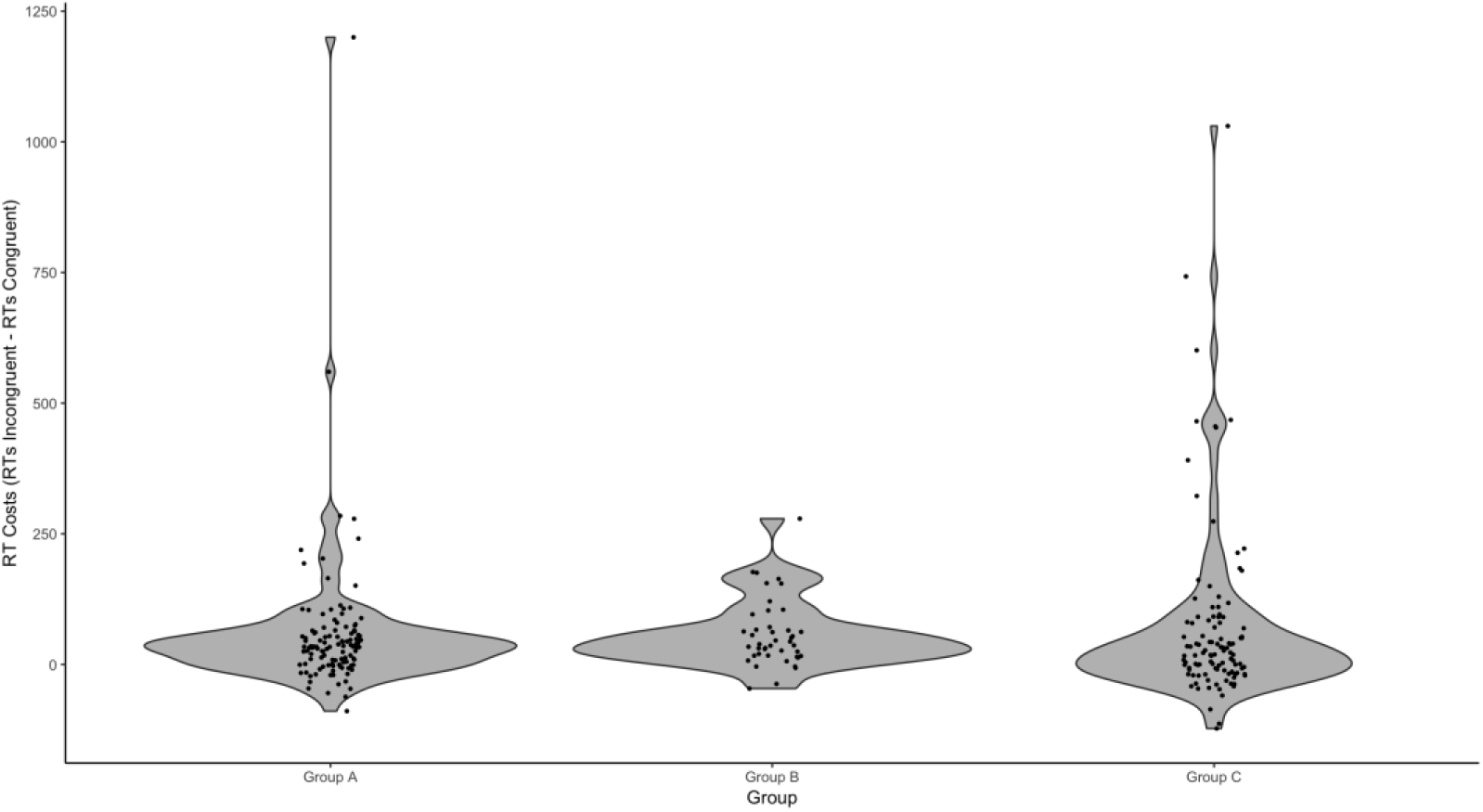
Distributions of RT difference between congruent and incongruent trials, for each group in Experiment 1.

### Discussion

The flanker effect was successfully replicated on a sample of 268 children tested using *Gorilla*. This characterised the ‘conflict network’, children taking longer to provide correct answers to incongruent trials, compared to congruent trials. This effect was lower than 100 ms (being 62.33 ms on average). As mentioned in the introduction, this small magnitude of difference requires both accurate visual timing and response recording to detect. Crucially, there was no interaction between the flanker effect and participants’ group, even if their testing conditions differed greatly: two groups were taken from schools, over a mobile phone internet connection, and the third group was taken from a University setting, over a communal internet connection. In the case of accuracy, we did find a group difference on running non-parametric tests, however it was shown that after accounting for the age difference between groups, this disappeared – which suggests this was not caused by the testing environment.

In each group, however, pupils were supervised by a trained experimenter who guided them through the task, and who checked the quality of the internet connection. One of the potential benefits of web-based research is in reaching participants in various places (e.g., their own house), allowing for broad and unsupervised testing. Therefore, Experiment 2 tested whether the flanker effect would hold under such conditions, recruiting adult participants over Prolific and without supervision.

## Experiment 2

### Methods

#### Participants

104 adults were recruited, five participants were excluded for not performing above chance (< 60% accuracy) in the task (these individuals also had an accuracy in excess of 3 standard deviations from the mean). This left a sample of 99 of adults (57.57% female), with a mean age of 30.32 (SD = 6.64), ranging from 19 to 40 years old.

All participants were recruited online, through the Prolific.ac website, which allows recruitment and administration of online tasks and questionnaires (Palan & Schitter, 2018). All participants were based in the United Kingdom and indicated normal or corrected to normal vision, English as a first language, and no history of mental illness or cognitive impairment. This experiment was conducted in line with Cauldron Science’s Ethics code – which complies with the Declaration of Helsinki (World Medical Association, 2013). Informed consent was obtained through an online form, participants were informed they could opt out during the experiment without loss of payment.

Compensation for the task was £0.60 GBP, which on average translated to a rate of £8.70 per hour, as participants took an average of 4 minutes and 8.36 seconds to complete the task.

In addition, the software recorded the operating system, web-browser and browser viewpoint size (the number of pixels that were displayed in the browser) of the users. The breakdown is shown in Table 3 and Table 4.

**Table 6:** Breakdown of Browsers and Operating Systems within the sample. Total percentages of the sample are included in brackets.

| Count (Percentage) |  |
| --- | --- |
| **Browser** |  |
| Chrome | 75 (75.76%) |
| Safari | 9 (9.09%) |
| Firefox | 9 (9.09%) |
| Edge | 3 (3.03%) |
| Other | 3 (3.03%) |
| **Operating System** |  |
| Windows 10 | 57 (57.58%) |
| Windows 7 | 17 (17.17%) |
| Mac OS 10 | 16 (16.16%) |
| Chromium | 5 (5.05%) |
| Windows 8 | 4 (4.04%) |

**Table 7:** Viewport characteristics of the adult sample’s web-browsers. The viewport is the area of a browser containing the information from a site.

|  | Mean (Pixels) | Std. Deviation (Pixels) | Range (Pixels) |
| --- | --- | --- | --- |
| **Horizontal** | 1496.13 | 218.85 | 1051 - 1920 |
| **Vertical** | 759.40 | 141.66 | 582 - 1266 |

#### Procedure

Participants completed the task on their own computers at home, and were not permitted to access the task on a tablet or smartphone. Before starting the task, participants read a description and instructions for taking part in the study, which asked them to open the experiment in a new window and note that the task would take around 5 minutes to complete (with an upper limit of 10 minutes). When the participants had consented to take part in the study on Prolific.ac, they were given a personalised link to the *Gorilla* website, where the experimental task was presented. First, a check was loaded to ensure they had not opened the task embedded in the Prolific website (an option that was available at the time of writing), which would minimise distraction. Then the main section was administered in the browser; on completion of this they returned to Prolific.ac with a link including a verification code to receive payment.

### Flanker Task

An adult version of the ‘conflict network’ flanker task, taken from the ANT was used Rueda et al. (2004). The mechanics, trial numbers and conditions of this task were identical to those used in Experiment 1; however, the stimuli were altered. The fish were replaced with arrows, as is typically done in adult studies (Fan et al., 2002; see Rueda et al., 2004) for a comparison of the child and adult versions). This is illustrated in Figure 6 and the time-course illustrated in Figure 7.

**Fig. 6:**
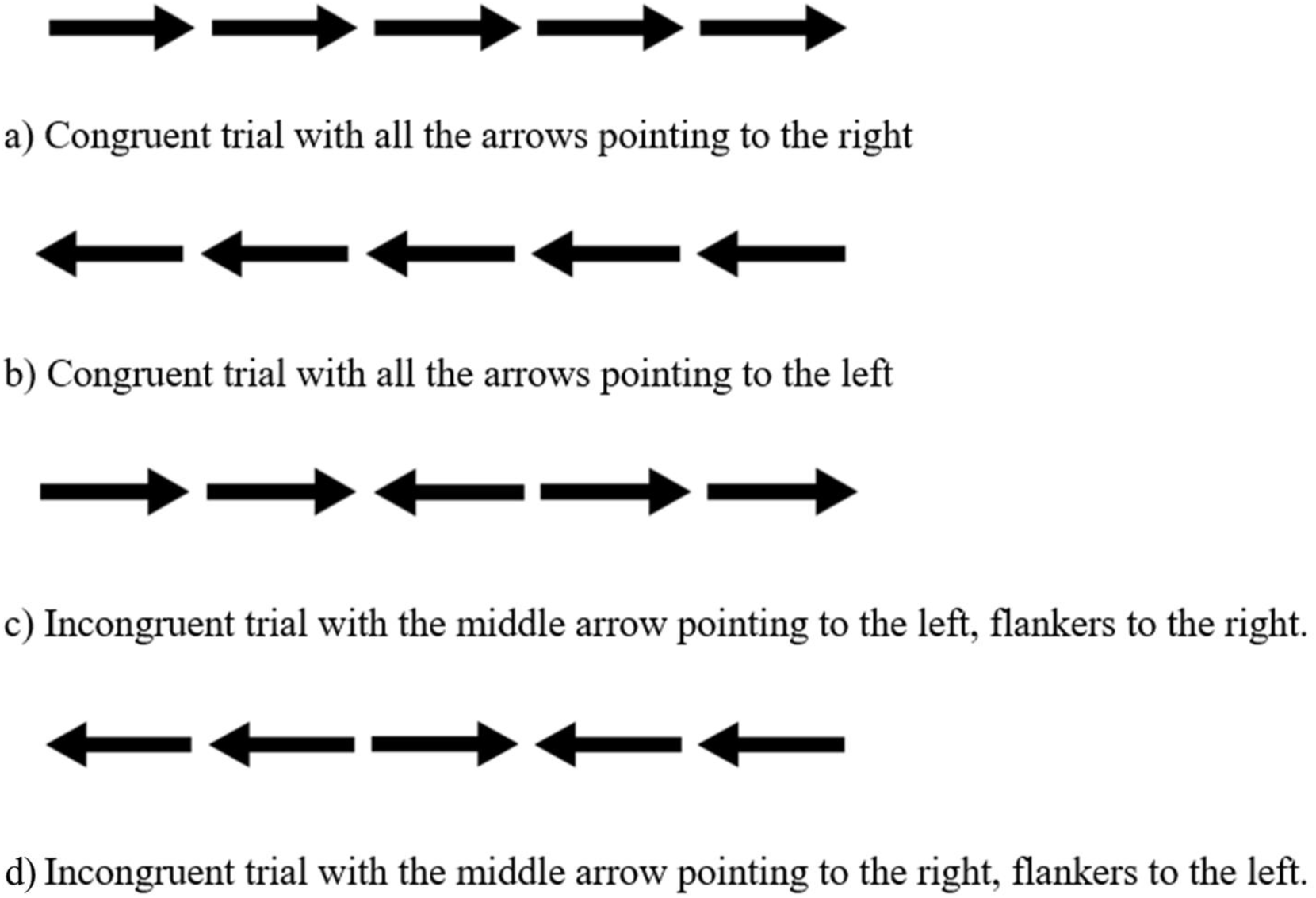
Trial types for Experiment 2. The different conditions used in the flanker task.

**Fig. 7:**
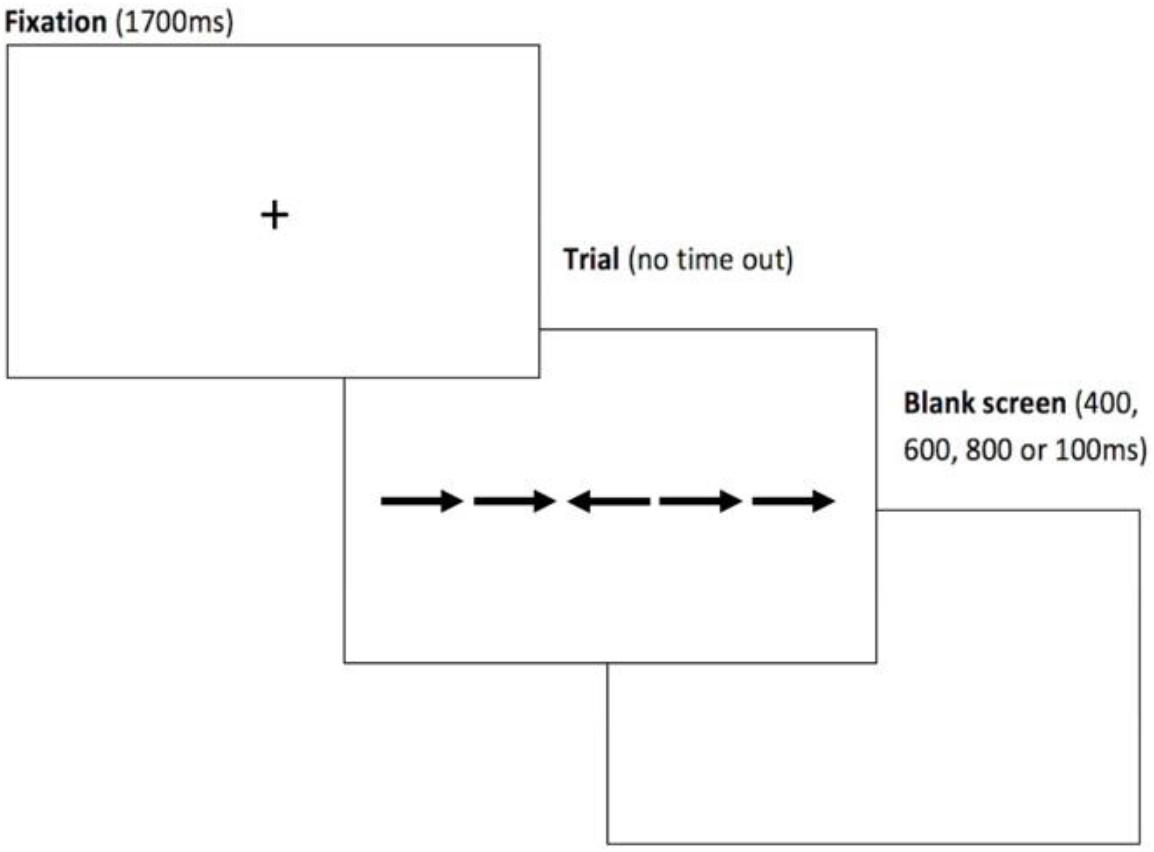
The time-course of a typical trial in Experiment 2. These screens represent what the participant was seeing within their web-browser.

Similarly to the children in Experiment 1, the adults were given written instructions, and then completed 12 practice trials with immediate feedback. Then moved on to complete 4 blocks of 24 trials (25% congruent-left, 25% congruent-right, 25% incongruent-left, 25% incongruent-right).

#### Power Calculations

The main effect of flanker reported in Rueda et al. (2004)’s adult arrow ANOVA results (Experiment 3) was *F*(2, 44) = 142.82; *p* = 0.0019. They did not report the effect size, so this permits us only to estimate the effect size using partial eta squared. This was calculated using the calculator provided by Lakens (2013), as *η*_p^2^_ = .87 (95% CI: .78 - .90).

However, as our planned comparisons for this group are simple (a t-test for mean RT and accuracy for incongruent versus congruent trials), we calculated power using the reported mean and standard deviation values from Fan et al. (2002) – Rueda et al. (2004) did not report the standard deviation, so this was not possible. The mean RT for congruent trials was 530 ms (SD = 49), and 605 ms (SD = 59) for incongruent trials. Using an a priori calculation from the G*Power software, this gave us a calculated effect size of d=1.38 and a sample size of 26 to reach a power of .96. However, this assumes that we are working in a comparable environment, which is not the case due to the increased potential noise - our sample size is therefore much larger than the original paper to account for increased noise, giving us a calculated power of <.99.

### Results

#### Data Pre-processing

As in Experiment 1, trials with RTs more than three standard deviations from the mean, and/or less than 200 ms were excluded from both accuracy and RT analyses.

#### Accuracy

Accuracy scores were computed over the total number of trials for each condition (congruent and incongruent). These means are shown in Table 5. As mentioned above, 5 participants were excluded for not being above chance based on these accuracy scores. Accuracy was distributed non-normally (Shapiro-Wilk W = 0.819 p <.001), so a Wilcoxon signed-rank test was used to compare the mean accuracy across the two types of trials. This provided evidence for a significant difference between the two means (1.72% difference, W = 1242, p < .001) with a Rank-Biserial Correlation of rrb=.49 (an estimation of effect size for non-parametric data, Hentschke and Stüttgen, 2011).

**Table 8.**
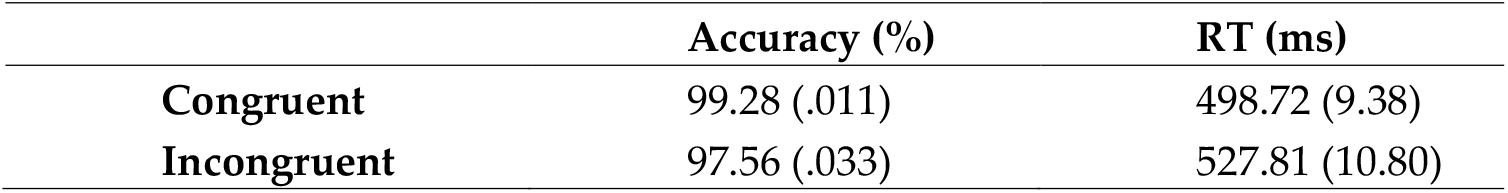
Average accuracy and correct trials reaction time for congruent and incongruent trials. Standard Errors are in brackets.

#### Reaction Time

Average RT was calculated for the two trial types - congruent and incongruent. Means and standard errors are reported in Table 5. RTs were only calculated for correct trials, as the accuracy rates were at ceiling. As above, a Shapiro-Wilk suggested the data was distributed non-normally (W = 0.748 *p* <.001), so a Wilcoxon signed-rank test was used to compare the differences in mean RTs. This test suggested a significant difference between the two means (29.1 ms difference, W = 414, *p* < .001) with a Rank-Biserial correlation of rrb=.83.

### Discussion

The ‘conflict network’ effect was observed and replicated. This was encouraging given the decrease in signal to noise that variance in operating system, web-browser, and screen size (shown above) would contribute towards this type of task. However, the effect of 29.1 ms was smaller than that observed in the original lab-based study (120 ms), and still smaller than the average effect of 109 ms reported in a meta-analysis of lab studies by MacLeod et al. (2010). This is likely due to variance in a remote environment, which may not be surprising, as MacLeod et al. (2010) found that there was large variance in the RT differences (congruent vs incongruent) between and within participants over multiple studies – 1655 ms and 305 ms respectively. Our smaller observed difference is also potentially driven by reduced RT variance. The average standard error in Experiment 1 was 20 ms, whereas it was around 10 ms in Experiment 2 – possibly leading to the lower than expected difference in RT. We are unable to compare our variance with the original paper’s child ANT results, as standard error or deviations were not reported. As a nearest online comparison, Crump et al. (2013)’s letter flanker’s difference between congruent and incongruent trials was 70 ms, which is closer to our observed difference, suggesting that online studies tend to find a smaller RT difference, however the stimuli and task structure differ significantly between our implementation and Crump et al. (2013)’s.

One potential explanation for the faster RTs, and decreased variance in the Prolific sample we tested could be their unique setting – the framing and task goals of these participants are different to typical volunteers. Research investigating users on the Mechanical Turk platform found that they were more attentive than panel participants (Hauser & Schwarz, 2016), suggesting internet populations are measurably different in their responses. Increased attentiveness could potentially lead to less within-subject variance – this may be an avenue of research for a future study.

## General Discussion

Gorrilla.sc is an Experiment Builder: a platform for the creation and administration of online behavioural experiments. It goes beyond an API, toolbox or JavaScript engine, and provides a full interface for task design and administration of experiments. It manages presentation time and response recording for the user, building on previous advances in browser-based research software without the requirement for programming or browser technology understanding. Utilising these tools, measurement of the ‘conflictnetwork’ was successfully replicated online. The replication persisted across several different groups, children in primary schools in two countries, children at a public engagement event, and adults taking part on their own machines at home. This demonstrates tasks built using this platform can be used in a wide range of situations – which have the potential to introduce unwanted variance in timing through software, hardware and internet connection speed – and still be robust enough to detect RT differences, even in a task containing a relatively low number of trials (< 100 trials).

Results such as these provide evidence that could enable more researchers to undertake behavioural research on the web, whilst also offering the maintained back-end which can be kept up-to-date with changes in user’s browsers - that otherwise would require a much higher level of technical involvement.

Building on these advantages, *Gorilla* is currently being used to teach research methods to undergraduate students in London at University College London and Birkbeck, University of London. In comparison with other software requiring specific programming skills, the teaching teams noted a lower need to provide technical assistance to students, allowing them to better focus on the research design per se.

### Limitations

Whilst technical involvement is lowered, there are still some limitations with presenting a task in the browser that the user should be aware of. These are mainly limited to timing issues, which *Gorilla* minimises but does not eliminate - there will always be an error rate, even though it is decreasing. The specific reasons for this error, and how it may be quantified or overcome in the future, are discussed below.

As with any software running on a user’s device, *Gorilla*’s response time is limited by the sampling/polling rate of input devices – a keyboard for example. Unfortunately, short of installing intrusive software on the user’s device, the web-browser has no mechanism for directly accessing polling rate - or controlling for polling rate. Often this sits at around 125 Hz, so this can be used to inform conclusions based on RT data gathered online. Future developments may at some point allow programs running in the browser to access hardware information and adjust for this - however, this will only be important for research which aims to model individual trials on an accuracy of less than 8 ms (the default USB polling rate for input devices is 125 Hz, so a sample every 8 ms). Alternatively, developments in recruitment platforms (such as Prolific and Mechanical Turk) may enable screening of participant’s hardware, allowing researchers to specify participants with high refresh monitors and high polling-rate input devices (most likely to be video-gamers). This would reduce the online research benefit of a larger available participant pool, but there are still many large and diverse groups of participants who meet such requirements, including the PC gaming community. Online research specifically targeting the gaming community has successfully gathered large amounts of data in the past (Ross, Irani, Silberman, Zaldivar & Tomlinson, 2010; Ipeirotis & Paritosh, 2011).

One unique problem in remote testing is the potential processing load any given participant may have running on their computer may vary dramatically. High processing loads will impact the consistency of stimulus presentation and the recording of responses. Fortunately, the platform records the actual time each frame is presented for, against the desired time - so the impact on timing can be recorded and monitored. A potential future tool would be a processing load check - this could either work by performing computations in the browser and timing them as a proxy for load. Or, it may potentially become possible to measure this using methods already available in Node.js (an off-browser JavaScript runtime engine) for profiling CPU performance – something that is likely to become possible if performance.now() timing is – at least partially – reinstated in browsers (for examples of how this could work see: Nakibly, Shelef, Yudilevich, 2015; Saito et al., 2016).

The use of modern browser features, such as requestAnimationFrame(), gives the best possible timing fidelity in the browser environment, and also allows for inconsistencies in frame refresh rate to be measured and accounted for. Online research will always be limited by the hardware that participants have, and despite the availability of modern monitors offering higher framerates, most users’ systems operate a refresh rate of 60Hz (Nakibly, Shelef & Yudilevich, 2015; Zotos & Herpers, 2012; 2016), therefore most stimulus presentation times are limited to multiples of 16.667 ms. Giving some insight into online participant’s device usage, an mTurk survey showed that over 60% were using either a laptop, phone or tablet – the vast majority of which will use a 60Hz refresh rate (Jacques & Kristensson, 2017). It is therefore advisable for users on any online platform to restrict presentation times to multiples of 16.667 ms. This is spoken about in *Gorilla*’s documentation, however, a future feature may be to include a warning to users when they try and enter non-multiples of the standard frame rate.

### Future Features

There are potential improvements to the platform that would make it a more powerful tool for researchers. These fall into two camps: tools for widening the range of experiments you can run, and tools for improving the quality of data you can collect.

In the authors’ experience, tools for researchers to run online visual perception, attention and cognition research are limited. This is perhaps a product of reluctance to use online methods, due to concerns regarding timing – which we hope to have moved towards addressing. In order to provide a greater range of tools a JavaScript-based Gabor patch generator is under development, which can be viewed using this link: www.bit.ly/GorillaGabor. This first asks participants to calibrate their presentation size to a credit card, and measure the distance to the screen - calculating visual degrees per pixel - and then allows presentation of a Gabor patch with size, frequency, window size in degrees. Experimenters can also set animations which change the phase and angle of these patches over time. These animations are fast (40Hz) as the patch and window are pre-generated and manipulated to produce the animation, rather than a frame-by-frame new patch generation.

Another tool that widens online research capabilities is remote, webcam-based eye tracking. An implementation of the WebGazer.js library (Papoutsaki et al., 2016) for eye-tracking is also being integrated into the platform. This permits rough eye-tracking, and head position tracking, using the user’s webcam. Recent research has provided evidence that this can be used for behavioural research, with reasonable accuracy - about 18% of screen-size (Semmelmann & Weigelt, 2018). This will also include a calibration tool, which can be run as frequently as needed, which allows the quantification of eye-tracking accuracy, and offers the ability to end the experiment if the webcam cannot be calibrated to the desired level. A prototype demo of the calibration is available here: www.bit.ly/EyeDemo. Additionally, WebGazer.js allows the experimenter to track the presence and changes in distance, of a user’s face. This can help with data quality, as you can assess when a user is looking at the screen, and prompt them to remain attentive to the task. The impact of this type of monitoring may be particularly interesting to investigate in a task such as the one presented in this paper – perhaps participants would show a different flanker effect if they were more attentive in the task.

## Conclusion

We described *Gorilla* as a tool which lowers the access barriers to running online experiments – e.g. understanding web development languages, servers, programming APIs – significantly, by managing all levels of implementation for the user and keeping up to date with changes in the browser ecosystem. We presented a case study, to demonstrate *Gorilla*’s capacity to be robust to environmental variance (from software, hardware and setting) during a timing task. An RT sensitive flanker effect – Rueda et al. (2004)’s ‘conflict network’ – is replicated in several populations and situations. There remain some constraints in running studies online - there may be future ways of tackling some of these constraints (i.e. specialist hardware). Future improvements to the platform include: a Gabor generator, webcam eye-tracking, and movement monitoring.

## Notes

## Acknowledgements

We would like to thank Edwin Dalmaijer, Nick Hodges, Marie Payne, Daniel C. Richardson, Hannah Spence, and, Will Webster for their feedback, and discussion of this paper.

## Open Practices Statement

None of the data reported here are available, as the participants – the majority of whom are children – did not consent for any data sharing. The experiments were not preregistered. Some materials (e.g. the flanker tasks) are available on www.gorilla.sc.

## Supplementary Material: A tutorial on constructing the Flanker task

The Flanker task was built in two steps. First, we created the Task. Secondly, we incorporated this Task in an Experiment, which allowed us to recruit participants.

### Creating the task

The Flanker task is a fairly simple task to program on Gorilla. Out of the five tabs of the Task Builder (Task Structure, Spreadsheet, Stimuli, Manipulations, Script), we only need to use to first three ones.

The “Task Structure” (highlighted in red below) tab allows us to define the different sections within the task, and the different screens within each section.

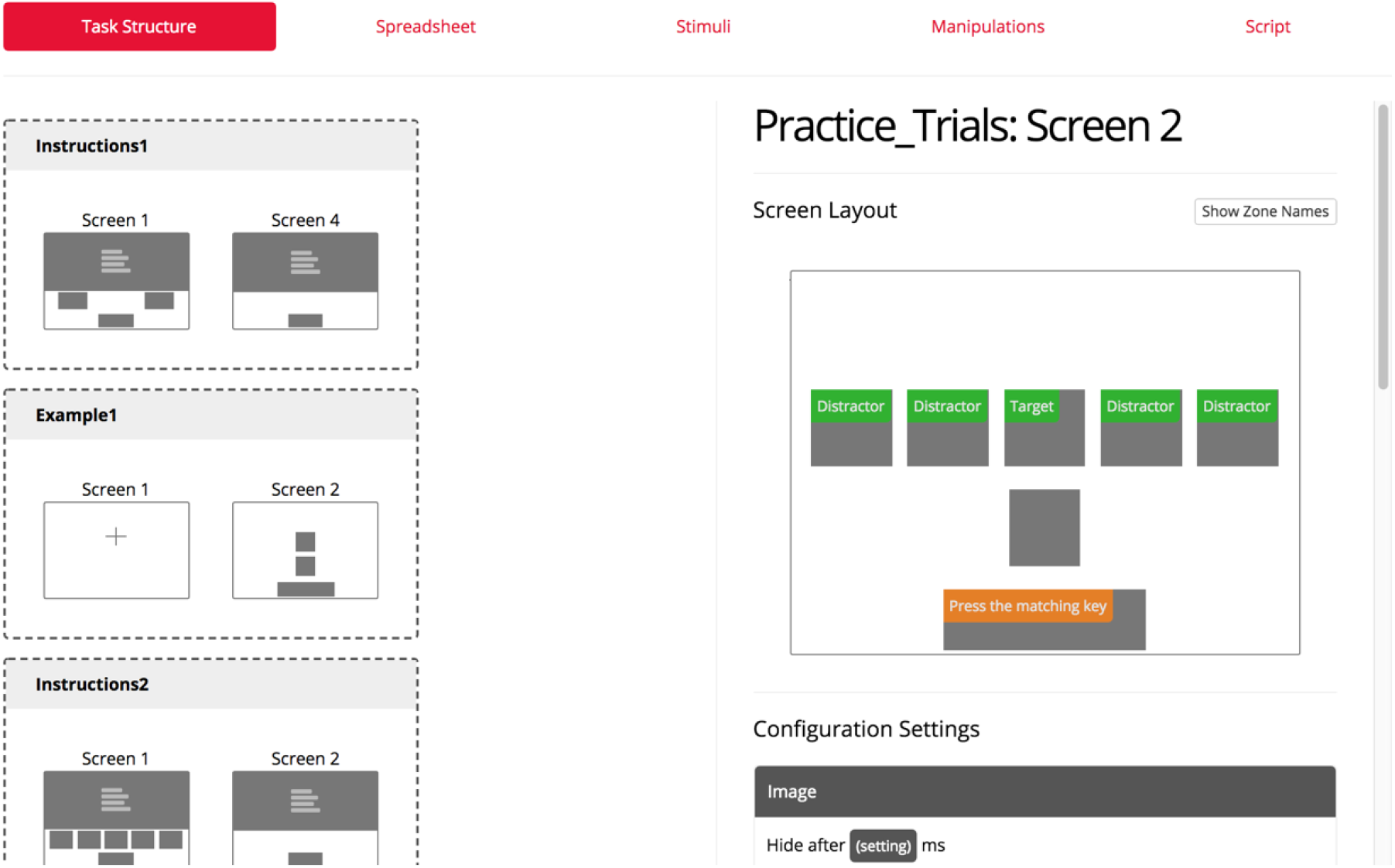

On the left, we can see the different sections. There is a first set of instructions (“Instructions 1”), followed by a very simple demonstration of the stimuli (“Example 1”). The instructions are fully developed in “Instructions 2” and followed by some practice trials (“Practice_Trials”).

On the right, we can see more specifically how the screen is designed for the practice trials. Five images zones have been defined. Their name is written in green: the central fish is the “Target”, the flanker fish the “Distractors”. The rectangular zone with an orange label indicates the response modality. Participants have to press a button saying whether the central fish is pointing to the right (“m”) or to the left (“z”). They will see the instructions that are written in orange on the screen: “Press the matching key”. The squared zone in between the green and orange zones is used to provide feedback. We will not develop it here since it is only used during practice, and not during the main task.

Crucially, the Screen set up only has to be defined once. The images that will populate it from trial to trial are specified in the Spreadsheet.

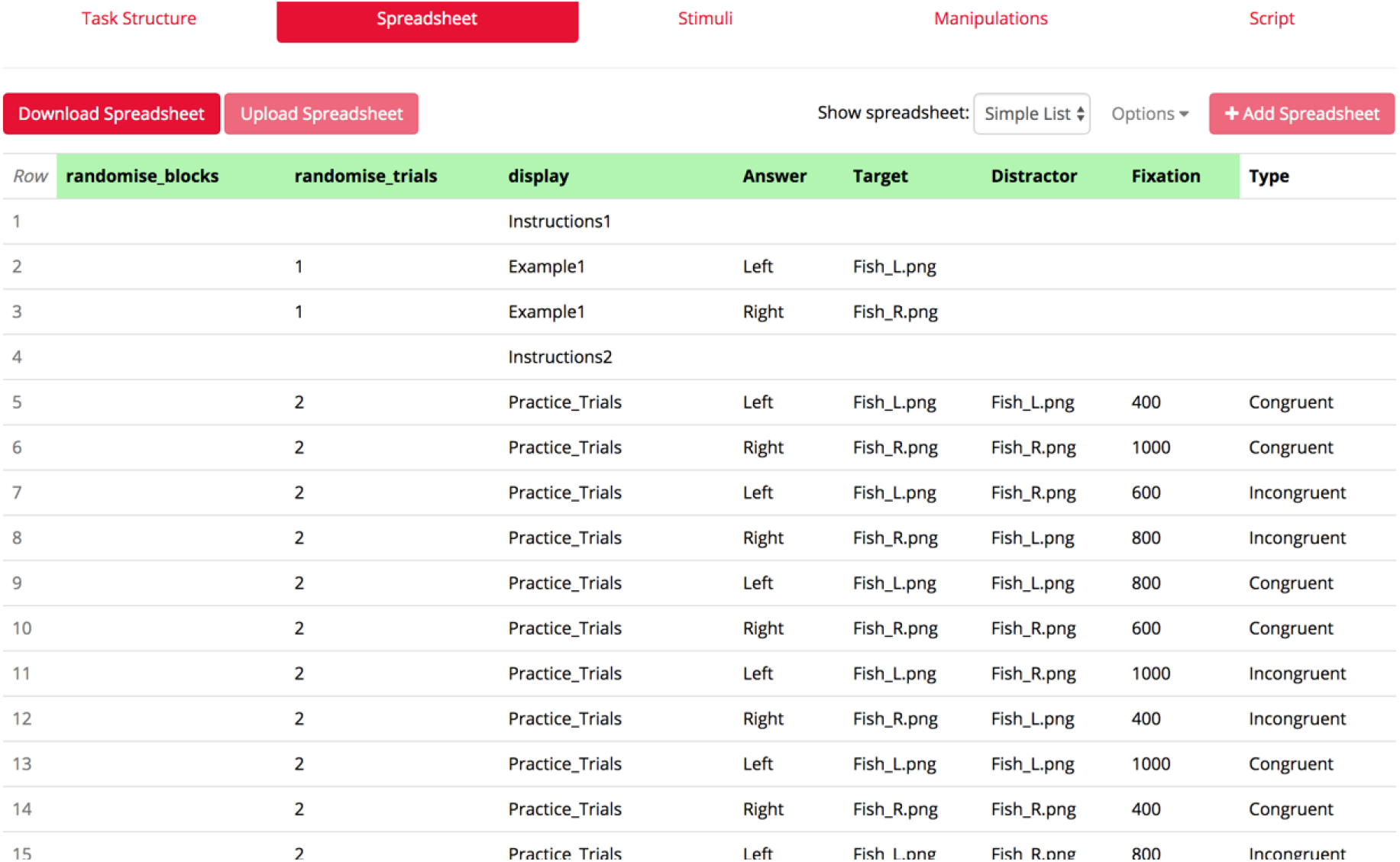

The “Display” columns mirrors the sections that have been created in our “Task Structure”. Let’s focus on the “Practice_Trials” display. In the “Target” and “Distractor” columns, we indicate which image will be uploaded for each trial. On the first Practice Trial, for example (Row 5), the Target is a fish pointing to the left, and the Distractors are also pointing to the left: it is actually a Congruent trial, as written in the Column “Type”. The images for each zone (here it is only “Fish_L.png”) have to be uploaded in the “Stimuli” menu.

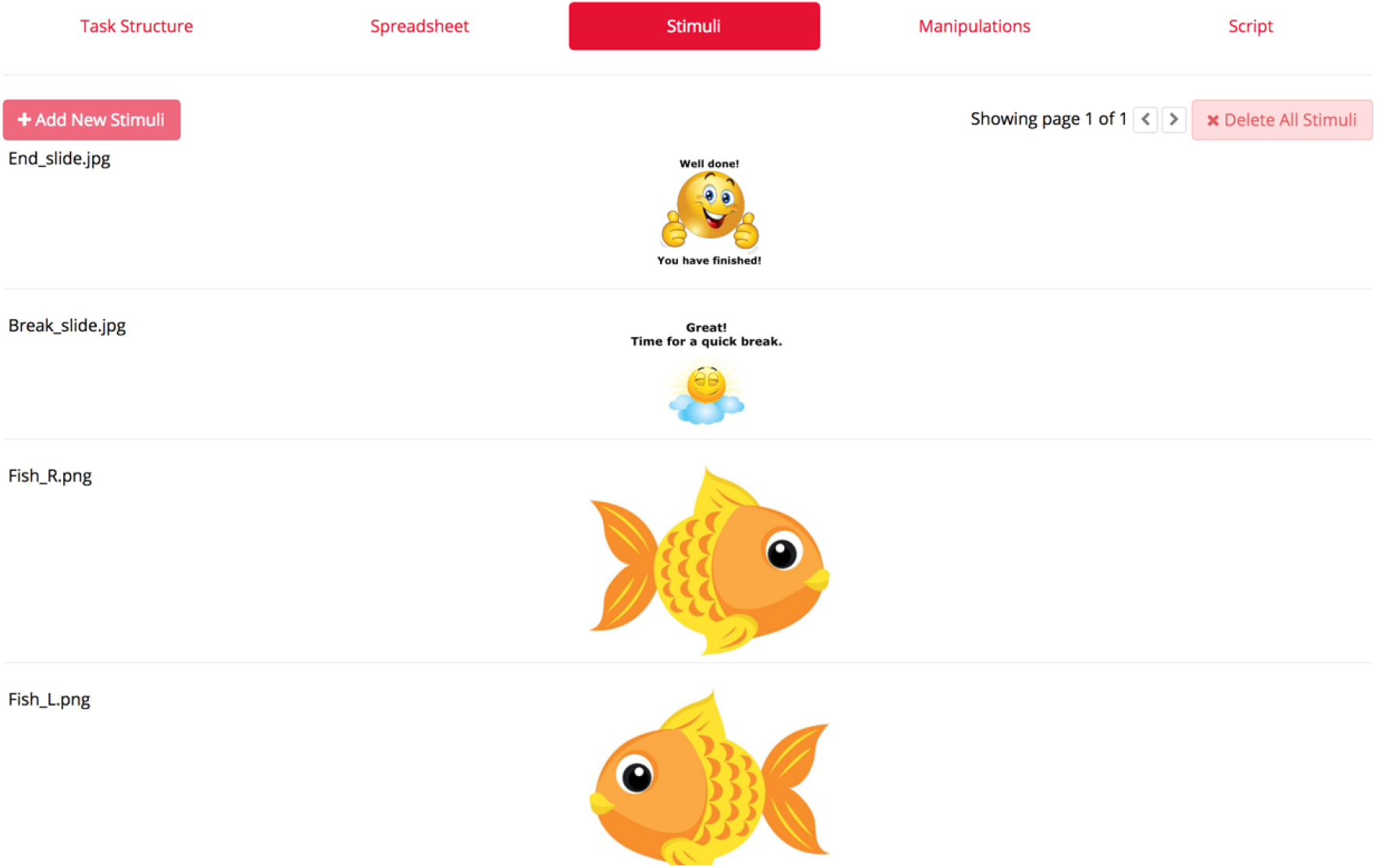

The last image is the one that is called “Fish_L.png”. Once we have set up our task, using the “Task structure”, the “Spreadsheet” and the “Stimuli” tabs, we need to put it in an experiment.

### Putting the task into an experiment

Gorilla uses what we call an “Experiment tree”. The tree specifies which tasks are used within an experiment, as well as their order. Here, the experiment (Called “Flanker Fish”) is pretty simple, because we only use the Flanker Task.

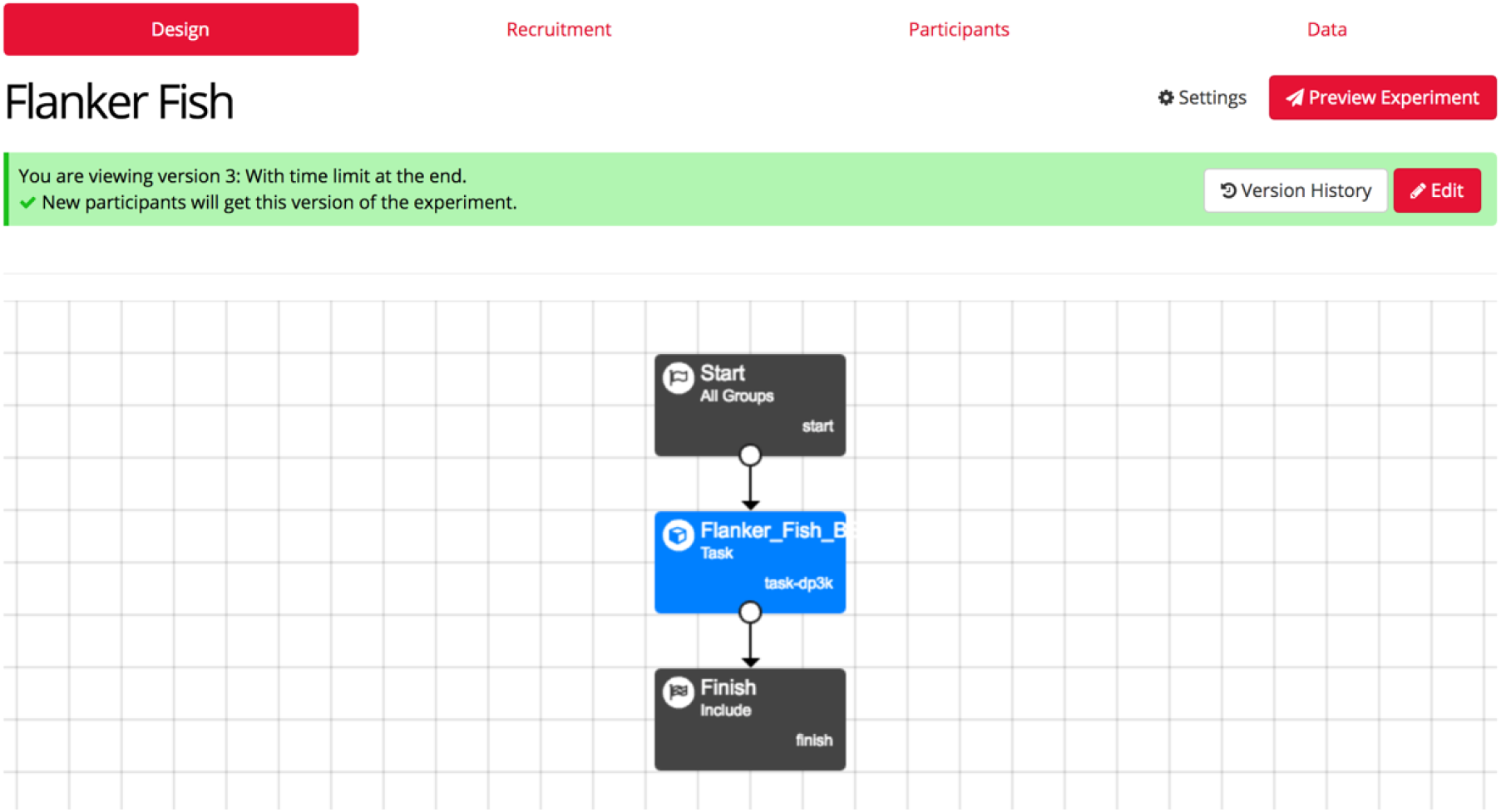

It is shown in blue, and it is surrounded by a “Start node”, and a “Finish node”, specifying the beginning and the end of the experiment. Many more functionalities are available in the “Experiment tree”, allowing researchers to counterbalance the order of presentation for an experiment that contains several tasks, for example.

The “Recruitment tab” allows us to generate a link to share the experiment online, and to select how many participants we would like to recruit, along with any specific technical requirement (device types, browser types of connection speed that the participants would need to have). The “Participants” tab references all the participants that joined the experiment, and the “Data” tab allows us to download our data.

## Footnotes

1 Note that recent developments in web-browsers have introduced different Application Programming Interfaces (APIs), for example the audio API in Chrome, allowing access to audio devices – calls can be made to these from JavaScript which are outside the event loop and are executed asynchronously. For a list of examples see: www.developer.mozilla.org/en-US/docs/Web/API

